# Dِistinct brain regions map olfactory and visual spaces

**DOI:** 10.1101/2025.09.11.675653

**Authors:** Heydar Davoudi, Jason R. Climer, John B. Issa, Daniel A. Dombeck

## Abstract

The hippocampus contains a multisensory cognitive map that incorporates spatial cues from across the senses, such as vision and olfaction. However, how primary sensory information transforms along pathways to the hippocampus into this single, combined map is largely unknown. Specifically, does the hippocampus generate or inherit the multisensory map of space? Here, we used multisensory virtual reality and large-scale functional imaging to determine if and how the main input regions to hippocampus, the lateral and medial entorhinal cortices (LEC and MEC, respectively), map olfactory and visual sensory spaces in navigating mice. We discovered that, in contrast to multisensory mapping in the hippocampus, LEC preferentially maps olfactory space while MEC preferentially maps visual space, regardless of the behavioral relevance of the spaces. This establishes the existence of largely independent brain maps for different sensory spaces, suggesting that the hippocampus builds the cognitive map by combining modality-specific maps from pre-synaptic cortical regions.

## INTRODUCTION

Navigation often requires animals to integrate cues from multiple sensory modalities into a cohesive and unified internal representation of space (Figure 1A)—a cognitive map <u>(Tolman 1948; Jacobs 2012; Bates and Wolbers 2014; Aboitiz and Montiel 2015; Epstein et al. 2017; Behrens et al. 2018; Whittington et al. 2022; Igarashi et al. 2022; Bruns and Röder 2023)</u>. The hippocampus plays the central role in this process <u>(O’Keefe and Dostrovsky 1971; O’Keefe and Nadel 1978; Wood et al. 1999; Aboitiz and Montiel 2015; S. Zhang and Manahan-Vaughan 2015; Fischler-Ruiz et al. 2021)</u>, as its place cells map multisensory spaces in a task-dependent manner <u>(O’Keefe and Dostrovsky 1971; Anderson and Jeffery 2003; Eichenbaum 2017; Radvansky et al. 2021; Zutshi et al. 2025)</u>. Yet, the mapping of different sensory modality spaces (vs. primary sensory encoding; Figure 1B) during navigation has not been systematically explored even one synapse upstream from the hippocampus–in the entorhinal cortex. Therefore, it is not yet known whether the hippocampus generates or inherits the multisensory map of space.

**Figure 1.**
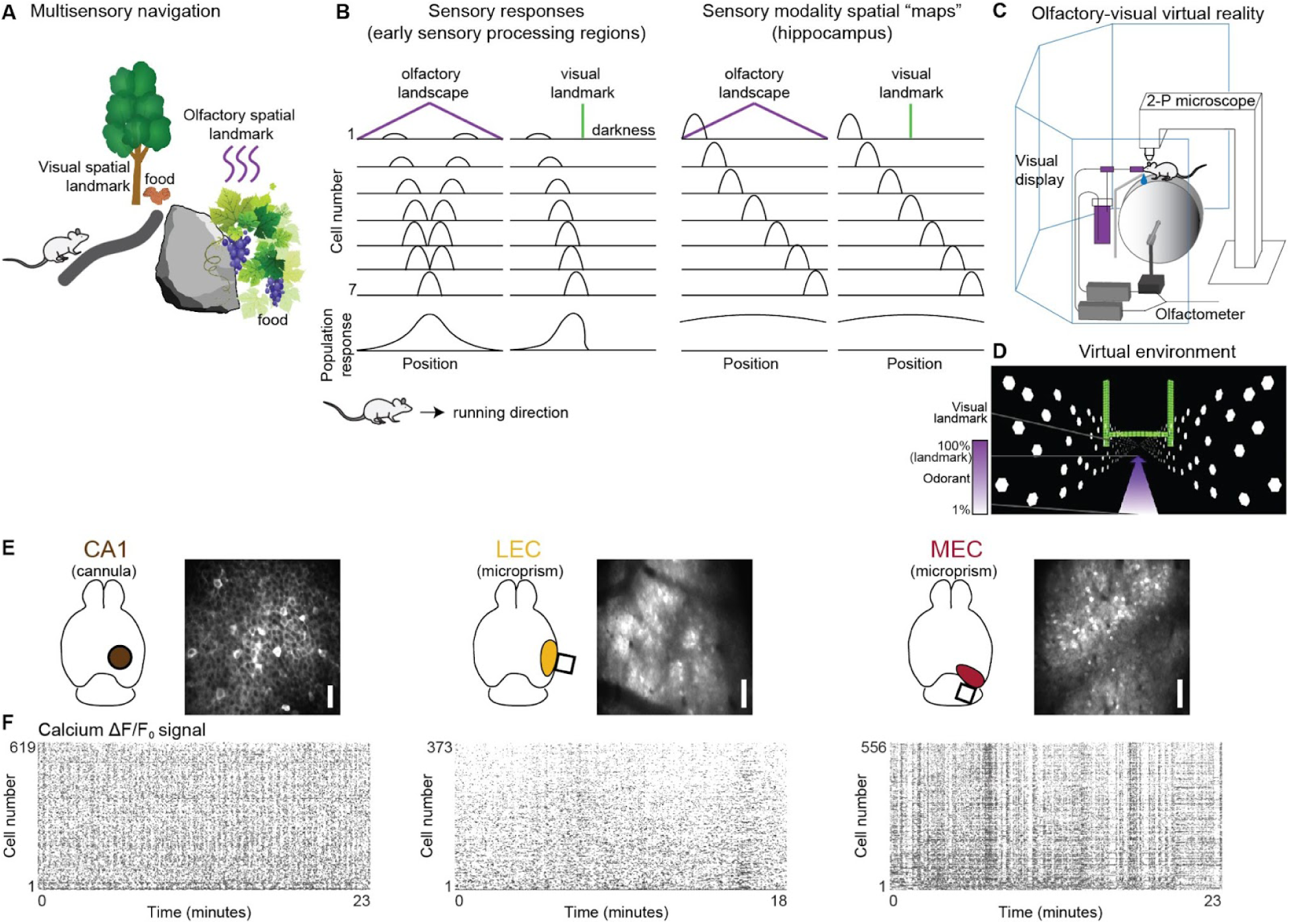
Entorhinal and hippocampal two-photon imaging during olfactory-visual virtual navigation. A) Schematic of a mouse in a multisensory habitat using visual and olfactory landmark-based navigation. Depending on environmental demands, animals may rely on visual landmarks, olfactory gradients, or a combination of both to find food. B) Schematic examples of how early sensory processing areas, such as olfactory bulb or retina/LGN (left), and cognitive brain regions (right) are likely to encode an environment defined by either a center peaked olfactory gradient (purple) or a single visual landmark (green) in a mouse running through the environment. In early sensory processing regions, encoding is expected to be driven directly by the properties of the sensory cues at each location. For example, in the olfactory bulb where neurons can respond to the identity and concentration of an odorant, a symmetric representation would be observed. In the retina, approaching a landmark would continuously increase the retinal space occupied, leading to an increasing response, which would then drop to zero once the object is passed and out of view. Cognitive brain regions, on the other hand, are expected to form a continuous map of these environments where every location is uniquely encoded, even different locations with the same sensory cues (as in the olfactory-defined environment) or with no sensory cues (as in the last half of the visually defined environment). C) The visual-olfactory multisensory experimental setup, including a head-fixed mouse running on a cylindrical treadmill surrounded by a five-panel VR display, an olfactory VR system ending with a nosecone over the mouse’s nose, and a water reward delivery system; all under a 2P microscope. D) In each traversal lap, the mouse experiences a visual landmark (green archway), visual optic flow on the walls, and an odor landmark defined as the peak of an odorant gradient (purple, not visible to mice). E) Left: 2-P imaging of dorsal CA1 neurons expressing GCaMP6m through a 2.5 mm diameter cranial window. Middle: 2-P microprism imaging of LEC neurons expressing GCaMP6s through a 2×2 mm microprism. Right: 2-P microprism imaging of MEC neurons expressing GCaMP6s through a 2×2 (or 1.5×1.5) mm microprism. White bar = 50 µm. F) ΔF/F₀ versus time traces for populations of CA1, LEC, and MEC neurons shown in (E).

The entorhinal cortex is anatomically divided into lateral (LEC) and medial (MEC) subdivisions, which differ in their anatomical inputs and presumed functional roles <u>(Witter et al. 2017; Nilssen et al. 2019)</u>. LEC receives a large fraction of its input from parahippocampal and olfactory areas <u>(Nilssen et al. 2019)</u>. During navigation, it has been shown to encode visual objects <u>(C. Wang et al. 2018; Deshmukh and Knierim 2011)</u> and rewards in visual virtual environments <u>(Issa et al. 2024; Jun et al. 2024; Bowler and Losonczy 2023)</u> and to be involved in odor associative learning in navigation <u>(Igarashi et al. 2014)</u> and non-navigation based tasks <u>(Li et al. 2017; Liu et al. 2023; Mena et al. 2025)</u>. Importantly, while these prior studies suggest that LEC might be able to form olfactory-visual multisensory spatial maps, none establish that LEC actually contains such a map of space during navigation. However, if it does form spatial maps, it is also possible that the maps would be olfactory-specific, since the previously described representations of other sensory modalities, such as vision, might instead relate to context or behavioral states <u>(Keene et al. 2016; Wilson, Watanabe, et al. 2013; Wilson, Langston, et al. 2013; Vandrey et al. 2020; Huang et al. 2023; Deshmukh and Knierim 2011; Issa et al. 2024)</u>. Thus, currently, evidence that LEC can map olfactory and/or visual spaces is lacking.

MEC, on the other hand, contains the well-known spatially-tuned grid cells <u>(Hafting et al. 2005; Høydal et al. 2019; Tukker et al. 2022)</u> as well as other spatial cells <u>(Sargolini et al. 2006; Solstad et al. 2008; Høydal et al. 2019)</u> in rodents during navigation of multisensory environments. The firing fields of grid cells were shown, in some conditions, to persist even in the dark <u>(Hafting et al. 2005)</u>, which could be interpreted to mean that grid cells map multisensory spaces, since they might be driven by sensory modalities other than vision. However, MEC is known to receive a large fraction of its input from visual areas <u>(Nilssen et al. 2019)</u>, and other studies under different conditions in the dark found largely degraded grid cell activity <u>(Pérez-Escobar et al. 2016; G. Chen et al. 2016; Waaga et al. 2022; Wen et al. 2024)</u>. So it is also possible that MEC preferentially maps spaces defined by visual cues <u>(Killian et al. 2012; Meister and Buffalo 2018; Wilming et al. 2018; Kinkhabwala et al. 2020; Tukker et al. 2022; G. Wang et al. 2025)</u>. Thus it remains an open question whether MEC can map both olfactory and visual spaces and participate in olfactory-visual multisensory spatial processing.

Overall, it remains unknown if and how the presynaptic regions to the hippocampus map different senses during multisensory navigation. To address this question, we employed a multisensory virtual reality (VR) system enabling precise and independent control of olfactory and visual spatial cues, combined with large-scale two-photon (2-P) functional imaging in CA1 of the hippocampus, LEC, and MEC regions of mice performing different navigation tasks (Figure 1C). We found that LEC preferentially maps olfactory space and MEC preferentially maps visual space, both with significantly less task-specific modulation compared to the hippocampus. These results uncover parallel modality-specific pathways containing relatively independent spatial maps, which could then be used to construct a unified, task-specific multisensory cognitive map in the hippocampus. Thus, while the hippocampus constructs a flexible, goal-directed spatial map for action planning <u>(Zutshi et al. 2025)</u>, the less flexible elemental spatial maps of individual sensory modalities are largely established presynaptically to the hippocampus within distinct subdivisions of the entorhinal cortex.

## RESULTS

In our experimental setup, mice navigated a multisensory virtual linear track that simultaneously provides two independent yet superimposed olfactory and visual spaces <u>(Radvansky and Dombeck 2018; Radvansky et al. 2021; Climer et al. 2025)</u>. Inspired by the mouse’s innate ability to employ a gradient ascent algorithm for odor plume tracking to reach an odor source <u>(Gire et al. 2016; Findley et al. 2021; Ackels et al. 2021; Baker et al. 2018)</u>, we designed an olfactory space based on a pine-scented linear odor gradient of constant slope, in which the “odor landmark” was defined by the location of the peak odorant concentration on the virtual linear track (Figure 1D). The visual space was defined entirely by a large salient green archway (the “visual landmark”) within two infinite white-dot walls providing optic flow. To determine whether CA1, LEC, and MEC map different sensory spaces and how the task relevance of the sensory modality impacts spatial mapping, we used three navigation tasks (olfactory-guided, visually-guided, and first-target guided) with identical olfactory-visual experience but different rewarding strategies. Depending on the respective task, the water-rewarded target was defined either as the olfactory landmark, visual landmark, or whichever landmark appeared first.

To establish the neural representation of visual, olfactory, and multisensory spaces during the different tasks, we recorded from neuronal populations in CA1, LEC, and MEC using two-photon calcium imaging (Figure 1E-F; 45,277 cells recorded from 15 mice across 156 sessions). CA1 imaging was performed through implanted cannula windows <u>(Dombeck et al. 2010)</u>, whereas LEC and MEC were accessed via implanted microprisms <u>(Heys et al. 2014; Issa et al. 2024)</u>. This experimental framework enabled us to determine if and how the entorhinal-hippocampal network maps different sensory modality spaces during multisensory navigation.

### LEC maps olfactory space

While a large amount of evidence demonstrates the multisensory nature of hippocampal maps <u>(Wood et al. 1999; McKenzie et al. 2014; Radvansky et al. 2021; Fischler-Ruiz et al. 2021; Zutshi et al. 2025)</u>, it remains unclear whether multisensory maps exist one synapse upstream to the hippocampus. To determine whether LEC and MEC map olfactory, visual or both sensory modality spaces, we first used the “olfactory-guided” navigation task, in which head-fixed mice received water rewards only at the odor target location of the multisensory linear track (Figure 2A). Therefore, in this paradigm, the olfactory space was behaviorally relevant and the visual space was not, serving as a distractor. On each trial, mice traversed the track in which they received a reward at the odor target, regardless of whether is was the first or second appearing landmark, then they experienced a brief pause at the second landmark (1 s freeze followed by 2 s without cues) before the next trial (STAR Methods). To ensure that mice relied on the sensory cues rather than idiothetic path integration (e.g., step counting; <u>(Pastalkova et al. 2008)</u>), the order and position of olfactory and visual landmarks and total track length were pseudo-randomized on each trial (Figure 2A). Track lengths ranged from 3.55 to 6.00 m (mean ± SD: 5.26 ± 0.6 m), landmark spacing from 1.15 to 3 m (2.01 ± 0.5 m), and the visual landmark appeared at one of seven possible positions (STAR Methods). Mice (n = 11; 52 sessions) showed strong task engagement, exhibiting anticipatory slowing and licking before the odor target (Figure 2B-C and S1A–B).

**Figure 2.**
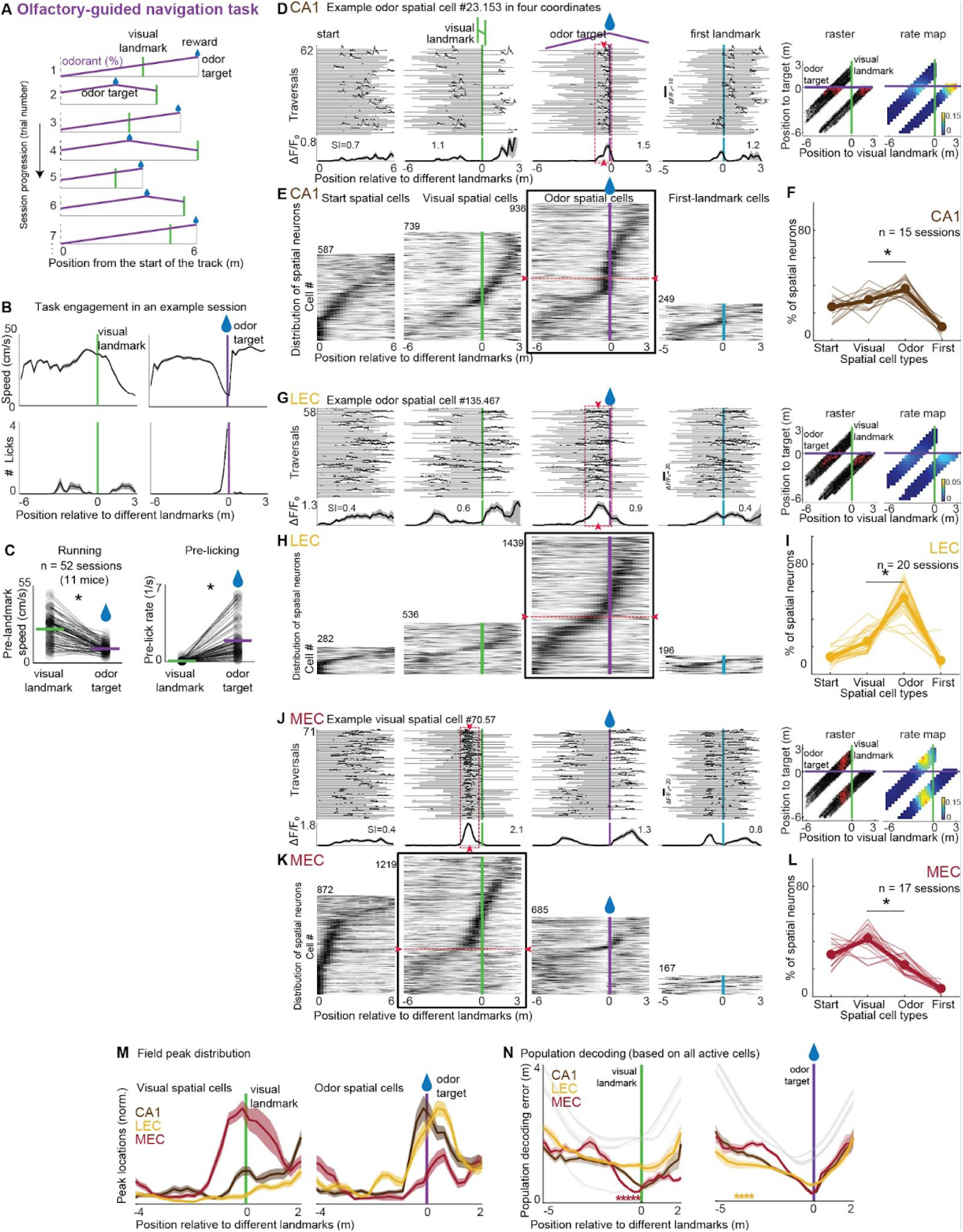
LEC maps olfactory space and MEC maps visual space during olfactory-guided navigation. A) The olfactory-guided navigation task. In each lap, mice experience a different odor target location (purple), visual landmark location (green bar), and track length and only receive water reward (blue droplet) at the odor target, where the odorant concentration gradient peaks. B-C) Task engagement quantified by anticipatory slowing (first row) and lick increase (second row; consumptive licks removed) relative to odor target (purple) but not visual landmark (green), as shown for an exemplar session (B). Task engagement for all sessions (C). 11 mice; n = 52 sessions; two-tailed Wilcoxon signed-rank test: speed, z(51) = 6.1, p = 1.0e-9; licking, z = –6.3, p = 3.7e-10. * p < 0.05. D) An example CA1 odor-spatial cell shown in four coordinates during an olfactory-guided navigation session. Top: Each horizontal trace represents the calcium trace (ΔF/F₀) of the cell in a traversal lap versus the position relative to four different landmarks, i.e., relative to the start of the track (first column), visual landmark (second target), rewarded (blue droplet) odor target (third column), and first landmark (fourth column). Bottom: The average spatial tuning of the example cell relative to four different landmarks. Spatial information (SI) value of spatial tuning in each coordinate is shown. The dashed red rectangle and arrowheads show the tuning field of the example odor-spatial cell. Right: for the example odor spatial cell, the raster plot of the MLE-deconvoluted spikes (red dots; STAR Methods) is depicted relative to locations of odor target and visual landmarks. These spikes are overlaid on the animal’s position (black dots). The rate map (in Hz) of the raster plot for the example cell is also shown. This example odor spatial cell is active before the odor target regardless of the visual landmark’s positions, resulting in a horizontal extent in raster and rate maps. E) All sorted and cross-validated CA1 spatial neurons from all sessions classified either as start-spatial, visual-spatial, odor-spatial, or first-landmark cells during the olfactory-guided navigation task. Each horizontal line shows the spatial tuning of each spatial neuron. The red arrowheads show the example odor-spatial cell in (D). The black box highlights the space most strongly mapped by CA1. F) The distribution of four CA1 spatial cell types during olfactory-guided navigation. Each trace shows the fraction of spatial cell types in a session (brown lines); the mean across sessions shown in thick brown. 3 mice, n = 15 sessions, two-tailed Wilcoxon signed-rank test: odor vs start, p = 8.5e-4; odor vs visual, p = 0.04; odor vs first, p = 6.1e-5. * p < 0.05. G-I) An example odor-spatial cell (G) and the populations of LEC spatial cells (H-I) as described above for CA1. 4 mice, 20 sessions, two-tailed Wilcoxon signed-rank test: odor vs start, p = 4.4e-4; odor vs visual, p = 4.4e-4; odor vs first, p = 4.4e-4. J-L) An example visual-spatial cell (J) and the populations of MEC spatial cells (K-L) as described above for CA1. 4 mice, 16 sessions, two-tailed Wilcoxon signed-rank test: visual vs start, p = 0.039; visual vs odor, p = 0.0013; visual vs first, p = 4.4e-4. M) Distribution of spatial field peaks of CA1, LEC, and MEC spatial cell populations. Visual (left) and odor (right) spatial cell peaks shown relative to visual and odor landmarks. N) Bayesian decoding errors of CA1, LEC, and MEC neural populations relative to the visual landmark location along the visual space (left) and odor target location along the olfactory space (right). * p < 0.05. See also Figure S1.

Next, to establish the mapping of visual and olfactory spaces during the olfactory-guided navigation task, we imaged neural populations within the hippocampal–entorhinal network. The task design allowed us to dissociate the neural mapping of visual and olfactory spaces based on each mouse’s position relative to the corresponding sensory landmarks. In addition to these sensory-defined spaces, the design also enabled analysis of position mapping relative to the track start, as well as abstract spatial mapping of the first landmark (i.e., the first cue encountered, regardless of modality; examined in more detail below). We first imaged the hippocampal CA1 region. For each active CA1 neuron, we calculated the mean significant calcium transients (ΔF/F₀) and the spatial information of its spatial tuning with respect to four different coordinates (i.e., track start, visual landmark, odor landmark, and modality-invariant first landmark; STAR methods; Figure 2D). Out of 5,164 active CA1 cells, 2,328 had significant spatial information in at least one coordinate (3 mice, 15 sessions, 5,164 active cells; 344.3 ± 20.1 (Mean ± SEM) active cells per session; 46.5 ± 3.3% spatial cells), and we defined each neuron’s preferred space as the one with highest significant spatial information (Figure 2D and S1C-D). This process identified four spatial cell types (i.e., start spatial cells, visual spatial cells, odor spatial cells, and first-landmark spatial cells), and we visualized the spatial tuning of these four populations in peak-sorted, cross-validated heatmaps (Figure 2E-F). This analysis revealed significantly more odor spatial cells than the other populations, though there were still a significant fraction of start and visual spatial cells and very few first target spatial cells (Figure 2F). The heat maps (Figure 2E), along with peak distribution histograms of the different CA1 spatial cell populations (Figure 2M) revealed mapping of the different spaces. In particular, there was robust mapping of the olfactory space with a larger fraction of the cells encoding the locations around the rewarded olfactory target (i.e., reward clustering). This spatial information-based analysis was based on single cell activity patterns. Therefore, we also used a separate population analysis method, i.e., Bayesian decoding using all active cells, to investigate how CA1 mapped the different spaces. This analysis provided a similar picture as the above spatial information-based analysis, confirming that CA1 carried information about mouse position relative to each of the landmarks, with the lowest decoding error relative (and near to) the odor target (Figure 2N). These results establish that CA1 encodes multiple spaces during the olfactory-guided task, with a preference for mapping the olfactory space. These findings largely are in line with our previous results of olfactory and visual space mapping in the hippocampus <u>(Radvansky et al. 2021)</u>.

In contrast to CA1, in LEC (4 mice, 20 sessions, 5,984 active cells; 299.2 ± 43.3 active cells per session; 37.6 ± 3.0% spatial cells) we found far more pronounced mapping of the olfactory space compared to the other spaces, with spatial cells primarily classified as odor-spatial cells during the olfactory-guided task (Figure 2G-I and S1C-D). The heat maps (Figure 2H), along with peak distribution histograms of the different LEC spatial cells populations (Figure 2M) revealed that the odor spatial cells tiled the full olfactory space. This pattern of tiling is a hallmark of spatial mapping, and is not consistent with early sensory processing since the same odor concentrations on either side of the odor landmark would result in symmetric cell tuning across the landmark (e.g. cells that are active both before and after the landmark, Figure 1B)–a pattern not observed here (see also Figure S1F). Similarly, Bayesian LEC population decoding revealed lower position decoding error across the olfactory space compared to the visual and other spaces (Figure 2N). These results establish the first evidence that LEC maps an external sensory space, and indicate that LEC preferentially maps odor, rather than visual, spaces in the olfactory-guided navigation task.

In contrast to both CA1 and LEC, in MEC (4 mice, 16 sessions, 4,121 active cells; 257.6 ± 21.7 active cells per session; 57.3 ± 3.26% spatial cells) we found more pronounced mapping of the (distractor) visual space compared to the other spaces, with spatial cells primarily classified as visual-spatial cells during the olfactory-guided task (Figure 2J-L and S1C–D). The heat maps (Figure 2K), along with peak distribution histograms of the different MEC spatial cells populations (Figure 2M) revealed that the visual spatial cells largely tiled the full visual space. This pattern of tiling is a hallmark of visual spatial mapping and is not consistent with early sensory encoding of the visual landmark, otherwise there would be little spatial tuning once the large archway is passed by the mouse–a pattern not observed here (Figure 1B, Figure 2K, M). Interestingly, MEC displayed more robust mapping of the track start space compared to LEC, consistent with its known role in path integration <u>(McNaughton et al. 2006; Campbell et al. 2021)</u>. Similarly, though MEC contained a population of odor-spatial cells, these cells did not appear to tile the full olfactory space as in LEC, and instead their information appeared to largely derive from decreased firing before the odor target (Figure S1G). This is more consistent with previously described speed cells in MEC <u>(Kropff et al. 2015; Hinman et al. 2016)</u> than olfactory space mapping cells. Bayesian MEC population decoding analysis revealed a similar story. While the lowest decoding error was observed close to both the visual landmark and odor target, a broader space around the visual landmark had low decoding error compared to the odor target (Figure 2N). These findings establish that MEC primarily maps visual space in the olfactory-guided navigation task, a remarkable contrast to LEC and CA1, particularly given that the visual space is not behaviorally relevant to the mice during this task.

Altogether, the above results highlight significant and previously unknown differences in how hippocampal and entorhinal cortical regions encode multisensory spaces. While CA1 mapped track start, odor and visual spaces, there was a preference for mapping the olfactory space. In contrast, LEC preferentially mapped odor, rather than visual, space and MEC preferentially mapped visual, rather than olfactory space, which can be observed by comparing the proportion of different spatial cell types across brain regions (Figure 2E, F, H, I, K, L). Similarly, through inspection of the differences in spatial cell tiling and decoding errors across regions (Figure 2M-N and S1E), it can be observed that the visual and olfactory spaces are most robustly encoded by MEC and LEC, respectively, with CA1 typically falling between the two. Moreover, population-level coding of each coordinate evolves along the navigational trajectory, such that as decoding accuracy for distance from the track start decreases (Figure S1E), decoding accuracy of olfactory and visual spaces increases in distinct brain regions (Figure 2N).

### LEC maps olfactory space and MEC visual space during visually-guided navigation

Our and others’ recent research established a representation of reward experience epochs in LEC <u>(Issa et al. 2024; Bowler and Losonczy 2023)</u>. Though the full tiling of the olfactory space by LEC described above for the olfactory-guided task is different from what is expected for a reward experience representation, it is likely that at least some elements of this LEC representation are related to reward experience rather than olfactory space mapping. Similarly, our recent research found differences in CA1 in encoding of odor and visual spaces depending on behavioral relevance <u>(Radvansky et al. 2021)</u>, and so it is unclear if or how the MEC and LEC populations might change based on the behavioral relevance of the different sensory modalities. Thus, we used the “visually-guided” navigation task to determine whether our above key results (from the olfactory-guided navigation task) are general or task-specific. All aspects of this task, including the superimposed visual and olfactory spaces, were identical to the olfactory-guided task except that the mice received water rewards only at the visual target location instead of the odor-target location (Figure 3A). This task design enabled us to examine if and how LEC and MEC map visual and olfactory spaces, and whether the behavioral relevance of the visual space (and irrelevance of the olfactory space) influences the mapping. Mice (14 mice, n = 65 sessions) robustly performed visually-guided navigation with high task engagement (Figure 3B-C and S2A–B).

**Figure 3.**
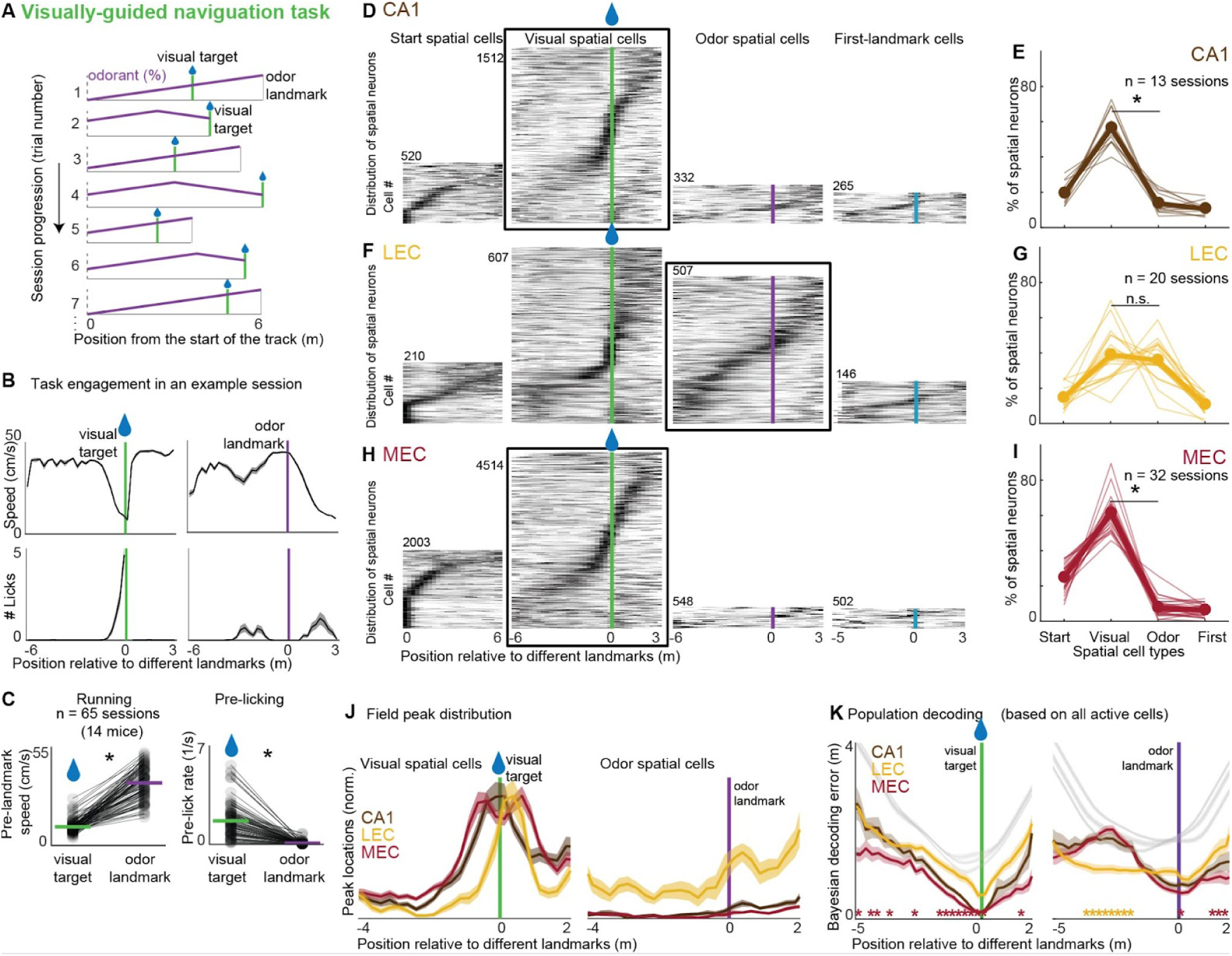
LEC maps olfactory space and MEC maps visual space during visually-guided navigation. A) The visually-guided navigation task. In each lap, mice experience a different odor landmark location, visual target location, and track length and only receive water reward at the visual target. B-C) Task engagement is quantified by anticipatory behaviors such as pre-reward slowing and increased licking prior to reaching the visual target, but not in response to the olfactory landmark for an exemplar session (B) and all sessions (C). 14 mice, n = 65 sessions; two-tailed paired Wilcoxon signed-rank test: speed, z(64) = –7.0, p = 2.4e-12; licking, z = 7.0, p = 2.4e-12. * p < 0.05. D-E) All cross-validated and sorted CA1 spatial cell types (D) and their distributions (E). The black box highlights the space most strongly mapped by CA1. Each brown trace shows the fraction of spatial cell types in a session (mean in thick brown trace). 3 mice, 13 sessions; two-tailed Wilcoxon signed-rank test: visual vs start, p = 2.4e-4; visual vs odor, p = 2.4e-4; visual vs first, p = 2.4e-4. F-G) All cross-validated and sorted LEC spatial cell types (F) and their distributions (G). 5 mice, 19 sessions; two-tailed Wilcoxon signed-rank test: odor vs start, p = 0.0012; odor vs visual, p = 0.77; odor vs first, p = 2.4e-4. H-I) All cross-validated and sorted MEC spatial cell types (H) and their distributions (I). 6 mice, 32 sessions; two-tailed Wilcoxon signed-rank test: visual vs start, p = 3.8e-6; visual vs odor, p = 3.8e-6; visual vs first, p = 3.8e-6. J) Distribution of spatial field peaks of CA1, LEC, and MEC spatial cell populations. Visual (left) and odor (right) spatial cell peaks respectively shown relative to visual and odor first landmarks. K) Bayesian decoding errors of CA1, LEC, and LEC neural populations relative to the visual target location along the visual space (left) and odor landmark location along the olfactory space (right). * p < 0.05. See also Figure S2.

We first imaged the hippocampal CA1 region in the visually-guided navigation task (3 mice, 13 sessions, 4,787 active cells; 368.2 ± 23.5 active cells per session; 48± 3.2% spatial cells) and found significantly more visual spatial cells than the other three populations, though there were still a small fraction of start spatial cells and very few odor spatial and first-landmark spatial cells (Figure 3D-E and S2C-D). The heat maps (Figure 3D), along with peak distribution histograms of the different CA1 spatial cell populations (Figure 3J) revealed robust mapping of the visual space and reward clustering around the visual target. Bayesian population analysis using all active cells provided a similar picture as the above analysis, confirming that CA1 carried information about mouse position in the visual space, with the lowest decoding error relative to (and near) the visual target (Figure 3K). These results establish that CA1 primarily and preferentially maps visual space during visually-guided multisensory navigation, consistent with our previous findings <u>(Radvansky et al. 2021)</u>.

In contrast to CA1, in LEC (5 mice, 19 sessions, 3,412 active cells; 179.6 ± 18.6 active cells per session; 33.7 ± 2.6% spatial cells) we found a pronounced mapping of the (distractor) olfactory space during the visually-guided task. We found that odor-spatial and visual-spatial cells were the primary spatial cell types, present in similar proportions (Figure 3F-G and S2C-D). The heat maps (Figure 3F), along with peak distribution histograms of the different LEC spatial cells populations (Figure 3J) revealed that LEC visual-spatial cells exhibited robust reward clustering around the visual target and firing fields before and after the reward reminiscent of our and others’ previously described reward experience epochs <u>(Issa et al. 2024; Bowler and Losonczy 2023)</u>. In contrast, the odor spatial cells tiled the full olfactory space in a way similar to that seen in the olfactory-guided task. Similarly, Bayesian LEC population decoding revealed relatively broad and homogeneously low decoding error across the olfactory space and low decoding error only at the rewarded visual target in visual space (Figure 3K). These results provide further support for our finding that LEC maps an external sensory space, and indicate that in the visually guided task, where the olfactory space was behaviorally irrelevant, LEC still preferentially mapped the olfactory space. At the same time, LEC neurons encoded aspects of reward experience near the visual target.

In contrast to LEC and similar to CA1, in MEC (6 mice, 32 sessions, 9,360 active cells; 292.5 ± 27.5 active cells per session; 62.1 ± 3.0% spatial cells) we found a pronounced mapping of the visual space compared to the other spaces during the visually-guided navigation task. We found that visual-spatial cells were the primary spatial cell type, with small fractions of cells assigned to the other three types (Figure 3H-I and S2C-D). The heat maps (Figure 3H), along with peak distribution histograms of the different MEC spatial cells populations (Figure 3J), revealed that the visual spatial cells largely tiled the full visual space with some mapping of the start space and little to no mapping of olfactory and first target spaces. Bayesian MEC population decoding analysis revealed significantly lower position decoding error across the visual space and around the rewarded visual target compared to the olfactory space (Figure 3K and S2). These findings establish that MEC primarily maps visual space during visually-guided navigation and provides further support for the idea of a visual space encoding preference in MEC.

Altogether, these results provide further support for our findings of significant and previously unknown differences in how hippocampal and entorhinal cortical regions encode multisensory spaces. As in the olfactory-guided navigation task, in the visually-guided task, MEC preferentially mapped the visual space, while LEC, in contrast, continued to map the olfactory space even though it was not behaviorally relevant. CA1 on the other hand preferentially mapped the visual space in the visually-guided task and the olfactory space in the olfactory-guided task. Thus, our results demonstrate significant task dependent mapping of the same multisensory environment in the hippocampal CA1, and preferential mapping of specific sensory modalities in MEC and LEC with significantly less dependence on the behavioral relevance of the modalities compared to hippocampus.

### CA1, rather than the entorhinal cortex, maps a modality-invariant space

So far, we established that during both olfactory-guided and visually-guided navigation tasks, LEC preferentially maps the olfactory space, MEC preferentially maps the visual space, and the downstream hippocampal CA1 preferentially maps the behaviorally relevant sensory space. However, rodents foraging for visible or odor-emitting hidden food might sometimes rely on an abstract, modality-invariant representation of distance to the nearest food, regardless of whether it is visible or odor-emitting. How the cortico-hippocampal network maps multisensory space, when spatial information from both modalities is required remains largely unknown. We have previously shown that a subset of CA1 neurons encode such a modality-invariant “first-target” space <u>(Radvansky et al. 2021)</u>. However, it remains unclear whether and how LEC and MEC contribute to encoding this abstract space. To determine whether LEC and MEC can map an abstract modality-invariant space that is dependent on olfactory and visual spatial sensory information, we used the “first-target guided” navigation task. All aspects of this task, including the superimposed visual and olfactory spaces, were identical to the olfactory-guided and visually-guided navigation tasks except that the mice received water rewards only at whichever (visual or odor) landmark appeared first along the track (Figure 4A). This task design not only enabled us to examine if and how LEC and MEC map the behaviorally-relevant abstract space but also to determine whether the modality preference of these regions persisted.

**Figure 4.**
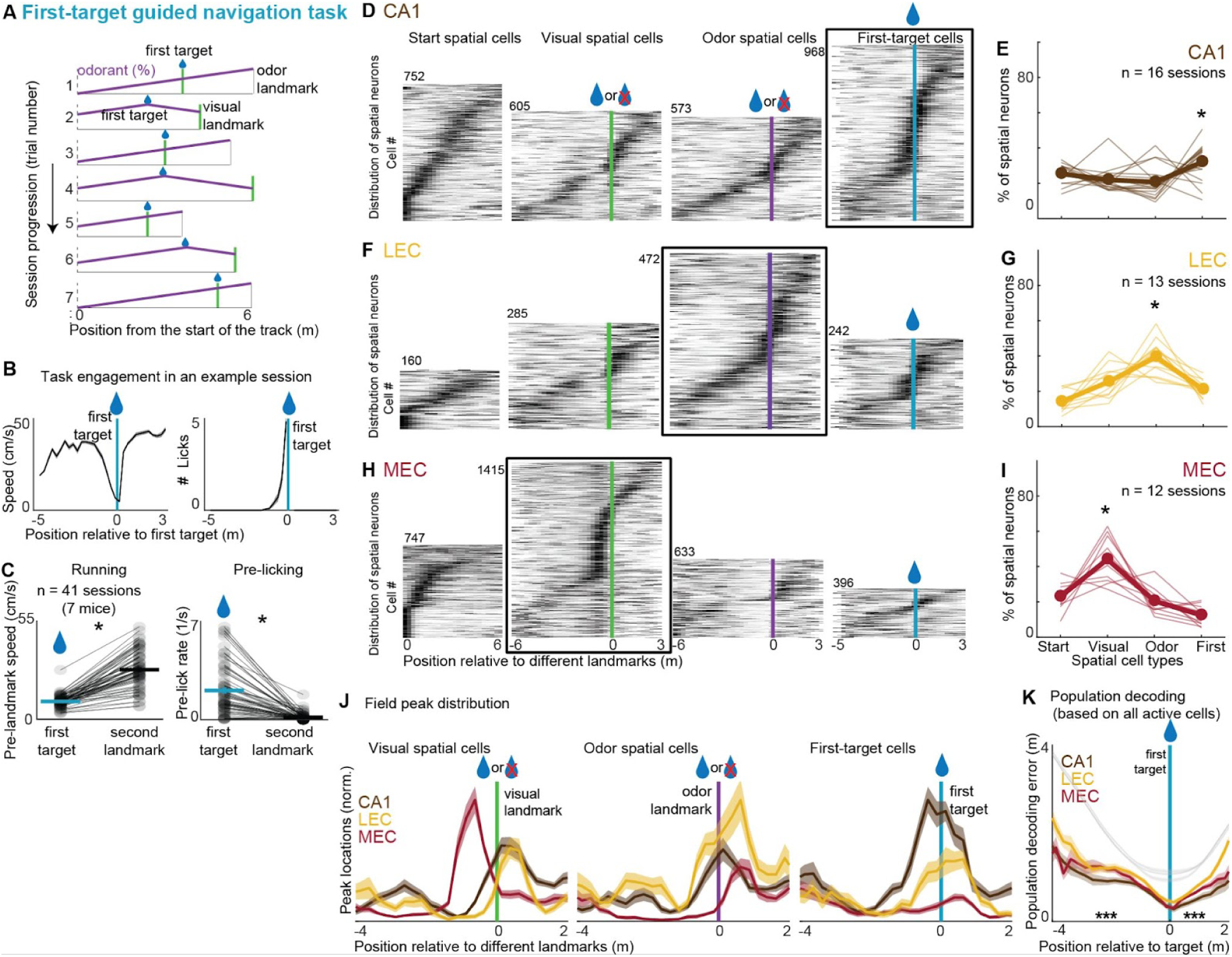
LEC maps olfactory space and MEC maps visual space during modality-invariant multisensory navigation. A) The first-target guided navigation task. In each lap, mice experience the same multisensory environment as in the olfactory-guided and visually-guided navigation tasks, with the only difference being that the reward location is at the first landmark, regardless of the landmark’s modality. |B-C) Task engagement is quantified by anticipatory behaviors such as pre-reward slowing and increased licking prior to reaching the first landmark, but not in response to the second landmark, irrespective of the landmark’s modality, as shown for an exemple session (B) and all sessions (C). 7 mice; n = 41 sessions; two-tailed Wilcoxon signed-rank test: speed, z(40) = –5.6, p = 2.4e-8; licking, z = 5.5, p = 4.1e-8. * p < 0.05. D-E) All cross-validated and sorted CA1 spatial cell types (D) and their distributions (E). While the first target is always rewarded (blue droplet) visual and olfactory landmarks may or may not be rewarded depending on their order of appearance in each traversal lap. The black box highlights the space most strongly mapped by CA1. Each thin grey trace shows the fraction of spatial cell types in a session (mean in thick trace). 3 mice, 16 sessions, two-tailed Wilcoxon signed-rank test: first vs start, p = 0.044; first vs visual, p = 0.039; first vs odor, p = 0.034. * p < 0.05 F-G) All cross-validated and sorted LEC spatial cell types (F) and their distributions (G). 2 mice, 13 sessions; two-tailed Wilcoxon signed-rank test: odor vs start, p = 9.8e-4; odor vs visual, p = 0.0049; odor vs first, p = 9.8e-4. H-I) All cross-validated and sorted MEC spatial cell types (H) and their distributions (I). 2 mice, 12 sessions, two-tailed Wilcoxon signed-rank test: visual vs start, p = 0.0024; visual vs odor, p = 9.8e-4; visual vs first, p = 4.9e-4. J) Distribution of spatial field peaks of CA1, LEC, and MEC spatial cell populations. Visual (left), odor (middle), and first-target (right) spatial cell peaks respectively shown relative to visual, odor, and first landmarks. K) Bayesian decoding errors of CA1, LEC, and MEC neural populations relative to the first target location. * p < 0.05. See also Figure S3.

Mice (7 mice, 41 sessions) robustly performed first-target guided navigation with high task engagement (Figure 4B-C and S3A–B). We imaged the hippocampal CA1 region in the first target task (3 mice, 16 sessions, 5,366 active cells; 335.4 ± 21.7 active cells per session; 50.5 ± 2.5% spatial cells) and the peak-sorted and cross-validated heat maps revealed significantly more modality invariant first-target spatial cells than the other populations, though there was still a significant fraction of start, visual spatial, and odor-spatial cells (Figure 4D-E and S3C-D). The heat maps (Figure 4D), along with peak distribution histograms of the different CA1 spatial cells populations (Figure 4J) revealed robust mapping of the first-target space and reward clustering around the first target. Through Bayesian population analysis of all active cells, we confirmed that CA1 carried significant information about mouse position in the abstract first-target space, with the lowest decoding error relative (and near to) the first landmark (Figure 4K). These results establish that CA1 preferentially maps the first-target space during first-target guided multisensory navigation, largely in line with our previous findings <u>(Radvansky et al. 2021)</u>.

In contrast to CA1, in LEC (2 mice, 13 sessions, 2,538 active cells; 195.2 ± 20.1 active cells per session; 34.4 ± 3.3% spatial cells) we found a pronounced mapping of the olfactory space during the first-target guided task. We found that odor-spatial cells were the primary spatial cell types that occurred, with a notable, though lesser, presence of start, visual-spatial, and first-landmark spatial cells (Figure 4F-G and S3C-D). The heat maps (Figure 4F), along with peak distribution histograms of the different LEC spatial cells populations (Figure 4J) revealed that LEC first target-spatial cells exhibited robust reward clustering around the first target and broad firing fields before and after the reward reminiscent of our and others’ previously described reward experience epochs <u>(Issa et al. 2024; Bowler and Losonczy 2023)</u> and similar to that observed in the visually-guided task. In contrast, the odor spatial cells tiled the full olfactory space in a way similar to that seen in the olfactory-guided and visually-guided navigation tasks. Importantly, when we separated trials where the olfactory landmark appeared first (i.e., the rewarded olfactory landmark) from those where it appeared second (i.e., the unrewarded olfactory landmark), we found that the odor spatial cells tiled the full olfactory space in both cases (Figure S3F). These results further establish LEC preferentially mapping of olfactory space, with evidence of this mapping now in a behavior paradigm that requires abstraction across sensory modalities.

In contrast to both CA1 and LEC, in MEC (2 mice, 12 sessions, 4,450 active cells; 370.8 ± 27.1 active cells per session; 54.3 ± 3.4% spatial cells) we found a pronounced mapping of the visual space during the first-target guided task. We found that visual-spatial cells were the primary spatial cell types that occurred, with a notable, though lesser, presence of start, odor-spatial, and first-target spatial cells (Figure 4H-I and S3C-D). The heat maps (Figure 4H), along with peak distribution histograms of the different MEC spatial cells populations (Figure 4J) revealed that the visual spatial cells tiled most of the visual space with more significant reward clustering compared to olfactory-guided and visually-guided tasks. Importantly, when we separated trials where the visual landmark appeared first (i.e., the rewarded visual landmark) from those where it appeared second (i.e., the unrewarded visual landmark), we found that the visual spatial cells tiled the full visual space in both cases (Figure S3F). MEC also displayed some mapping of the track start, olfactory and first target spaces. These results further establish MEC preferentially mapping of visual space, with evidence of this mapping now in a behavior paradigm that requires abstraction across sensory modalities.

These findings from the first-target guided navigation task are further supported by population-level analyses (Figure 4J-K, S3E-F). As demonstrated above, while LEC and MEC exhibited robust mapping of the odor and visual spaces, respectively, CA1 showed a markedly dense mapping of the abstract first-target space (Figure 4J). Additionally, CA1 exhibited the lowest Bayesian population decoding error across the first-target space (Figure 4K). Thus, as in the olfactory-guided and visually-guided navigation tasks, in the first-target task, LEC preferentially mapped the olfactory space, while MEC, in contrast, preferentially mapped the visual space. CA1, on the other hand, preferentially mapped the olfactory space in the olfactory-guided task, the visual space in the visually-guided task and the modality-invariant first target space in the first-target guided task.

### Mapping is sensory modality-specific in the entorhinal cortex and task-specific in CA1

As a summary of, and to further quantify, our findings, we directly compared the proportion of different cell types across brain regions and tasks (Figure 5A). In the olfactory-guided navigation task, LEC and CA1 preferentially mapped the rewarded olfactory space while MEC preferentially mapped the visual space. In the visually-guided navigation task, CA1 and MEC strongly mapped the visual space, while LEC continued to map the olfactory space, but with reward clustering around the visual target. During the first-target guided navigation task, while MEC preferentially mapped the visual space and LEC preferentially mapped the olfactory space, CA1 preferentially mapped the first-target space (Figure 5A; Table S1). Importantly, across the brain regions we found strikingly different levels of task modulations, as reflected in changes in the proportions of cells mapping the olfactory, visual, and first-target spaces across tasks (Figure 5B; Table S1). CA1 demonstrated dramatic task modulation: its neural population primarily mapped the behaviorally relevant space, i.e., olfactory space in the olfactory-guided task, visual space in the visually-guided task, and the abstract first-target space in the first-target guided task. In contrast to CA1, LEC and MEC displayed far less task modulation, with LEC and MEC preserving their preference for mapping the olfactory and visual spaces across the three tasks, respectively, with only some modulations in the fractions of cell types based on the behavioral relevance of the space (Figure 5B).

**Figure 5.**
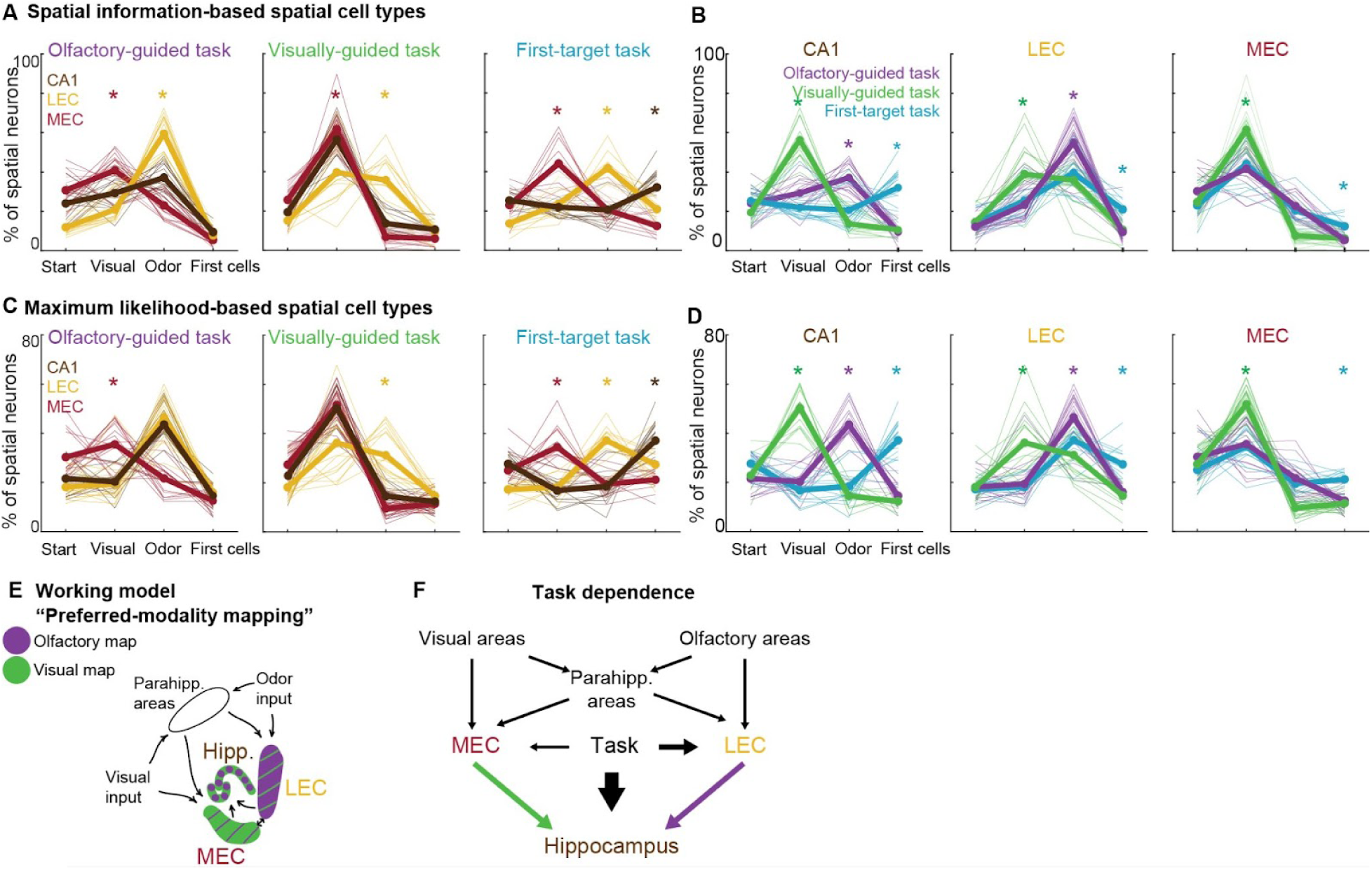
Mapping is sensory modality-specific in the entorhinal cortex and task-specific in CA1. A) The distribution of four spatial-information-based spatial cell types derived from CA1, LEC, and MEC during olfactory-guided (left), visually-guided (middle), and first-target guided (right) navigation tasks. Each thin trace shows the fraction of spatial cell types in a session (thick lines = mean). The asterisks and their colors indicate cases where the fraction of a particular cell type is statistically higher (p < 0.05) in one brain region compared to the other two brain regions. Detailed statistics are reported in Table S1. B) The distribution of four spatial-information-based spatial cell types in the three navigation tasks in CA1 (left), LEC (middle), and MEC (right). Each thin trace shows the fraction of spatial cell types in a session (thick lines = mean). The asterisks and their colors indicate cases where the fraction of a particular cell type is statistically higher (p < 0.05) in one task compared to the other two tasks. Detailed statistics are reported in Table S1. C) The percentage of MLE-based start, visual, odor, and first-target spatial cells in olfactory-guided (left), visually-guided (middle), and first-target guided (right) tasks. Each thin trace shows a session, with different colors showing different brain regions, and thick colors show the averages of sessions. Detailed statistics are reported in Table S1. D) The percentage of the same four MLE-based spatial cell types across tasks for each brain region, including CA1 (left), LEC (middle), and MEC (right) recorded during different multisensory tasks. Detailed statistics are reported in Table S1. E) Schematics of the proposed multisensory spatial representation model in the hippocampal-entorhinal network. F) A general schematic of spatial sensory pathways and task modulations in the hippocampal-entorhinal network. See also Figure S4 and S5.

Our findings so far have relied mostly on our spatial information-based classification of functional cell types (Figure 5A-B) and Bayesian population analysis (Figure 2N, 3N, 4N). To further verify our findings, we used a third independent method, the maximum likelihood estimation (MLE) framework (STAR Methods). We chose this method because our spatial information-based method does not consider conjunctive spatial tunings of individual cells, for example a cell with significant information in both visual and olfactory reference frames, but the MLE framework allows us to quantify any such conjunctivity. For this analysis, we quantified the cross-validated goodness-of-fit of each cell’s response in all possible spaces, including conjunctive spaces between two spatial coordinates (Figure S4A-B, S5; STAR Methods). Of the total 45,277 recorded active cells across different brain regions and tasks, 21,909 (48.8%) were identified as spatial cells using our MLE approach, of which 84.2% were classified as pure (non-conjunctive) start, visual, odor, first-target spatial cell types (Figure S4C). We examined the proportion of these MLE-identified, four non-conjunctive cell type populations and found that their distributions across brain regions and navigation tasks were highly similar to those revealed by our spatial information-based classification (Figure 5C-D; Table S1). We did, however, identify some conjunctive spatial processing across different brain areas and tasks, with these spatial cells (i.e., the remaining 15.8% of spatial cells) identified as conjunctive start-visual, start-odor, start-first, and visual-odor spatial cells; these cells are described in more detail in supplemental data (Figure S4D-G and S5). Thus, even with an MLE analysis approach capable of examining conjunctivity, the vast majority of active cells predominantly encoded one of our four previously defined spaces (i.e, start, visual, odor, or first-target) and, importantly, the distribution of these cells across brain regions and tasks confirms our above key results.

We therefore established, using multiple independent analysis methods, that behavior significantly modulates the mapping of multisensory space in CA1, but the entorhinal cortex largely maintains preferential and modality specific mapping despite changes in behavior. Thus, we propose a new working model of multisensory mapping in the entorhinal cortex, the “preferred-modality mapping” model, in which MEC and LEC predominantly represent visual and olfactory spaces, respectively, with a smaller fraction of cells in each region responding to the other sensory modality and significantly less behavioral modulation compared to CA1 (Figure 5E, F). In this framework, CA1 may use modality-preferred spatial maps from its upstream entorhinal inputs, to perform flexible, and if required, modality-invariant computations necessary for multisensory navigation.

## DISCUSSION

While olfaction is traditionally considered a contextual cue due to the perception that it is slowly changing or carries little spatial information, many studies have found a role for olfactory-guided navigation across many species <u>(Steck et al. 2010; Aboitiz and Montiel 2015; Gire et al. 2016; Hoy et al. 2016; Findley et al. 2021; Marin et al. 2021)</u>. Indeed, the sense of olfaction contains significant spatial information, comparable in many cases to the sense of vision <u>(McKissick et al. 2024; Barwich 2025)</u>, and also it has been shown that animals are capable of rapid, sub-sniff rate odor detection to support olfactory-guided navigation <u>(Ackels 2022)</u>. While place-related activity has been observed in primary olfactory areas such as the olfactory bulb <u>(Sterrett et al. 2025)</u> and the piriform cortex <u>(Poo et al. 2022; Mena et al. 2025)</u> in freely moving animals, a brain region that maps the full olfactory space of an animal’s local environment during navigation had not yet been identified. Our findings establish LEC as such a region that robustly maps the olfactory space across different behavioral contexts. LEC receives direct input from both the olfactory bulb <u>(Gretenkord et al. 2019; Sosulski et al. 2011)</u> and the piriform cortex <u>(Leitner et al. 2016; Bitzenhofer et al. 2022)</u>, and is known to be modulated by these regions <u>(Salimi et al. 2021; 2022; Y.-N. Chen et al. 2023)</u>. We speculate that primary olfactory sensory inputs are conveyed from these regions to LEC, where such information may be combined with self-motion signals to form the spatial map of olfactory space that we observed. Components of this spatial map may then be transmitted from LEC back to the olfactory bulb and piriform cortex, accounting for the previous observations of place-related activity in those regions. Notably, this olfactory mapping represents the first clear evidence of any spatial map in LEC. LEC neurons have previously been shown to encode non-spatial or egocentric spatial information <u>(Deshmukh and Knierim 2011; Wilson, Langston, et al. 2013; Wilson, Watanabe, et al. 2013; Keene et al. 2016; C. Wang et al. 2018; Vandrey et al. 2020; Huang et al. 2023)</u>, and they have been implicated in processing reward <u>(Issa et al. 2024; Bowler and Losonczy 2023; Jun et al. 2024)</u>. Our results add spatial mapping of olfactory spaces to this list for LEC and lead to the question of how one region encodes so many different variables related to behavior. One possibility is that LEC consists of many distinct, parallel circuits, each supporting the encoding of different representational domains. Indeed here, in addition to a map of olfactory space, we also observed reward-related encoding, consistent with this idea of parallel circuits in LEC.

MEC, on the other hand, has been extensively implicated in visual spatial coding (Hafting et al. 2005; Sargolini et al. 2006; Solstad et al. 2008; Killian et al. 2012; Heys et al. 2014; Aronov and Tank 2014; Low et al. 2014; Kropff et al. 2015; Meister and Buffalo 2018; Wilming et al. 2018; Høydal et al. 2019; Kinkhabwala et al. 2020; Tukker et al. 2022), visual cue processing (G. Wang et al. 2025), as well as in encoding reward (Butler et al. 2019; Boccara et al. 2019; Grienberger and Magee 2022), time (Heys and Dombeck 2018; Heys et al. 2020), and cognitive spaces such as sound (Aronov et al. 2017; Nguyen et al. 2024). Recent studies have demonstrated audiovisual sensory integration in MEC (Nguyen et al. 2024), and shown that MEC activity is more disrupted by visual than tactile deprivation (Tian et al. 2024). However, the capacity of MEC to map olfactory space, or to perform olfactory-visual spatial integration, remained unknown. Here, we found that MEC consistently mapped visual rather than olfactory space, even when vision was not behaviorally relevant, and with only minor task-related modulation. A small subset of MEC neurons in the olfactory-guided navigation task were classified as odor spatial cells, but closer inspection suggests that these were largely speed cells or speed-modulated grid cells (Figure S1D,G) (Kropff et al. 2015; Hinman et al. 2016), whose activity decreased as animals slowed down before reaching the rewarded odor target. While grid cells in MEC have traditionally been assumed to map multisensory spaces (Hafting et al. 2005; Høydal et al. 2019; Tukker et al. 2022), our results bring up the intriguing possibility that grid cells may predominantly map spaces defined by visual landmarks. However, our findings here come from spatially tuned neurons in 1-dimensional tracks (not necessarily grid cells) and so more research will be required to more clearly determine the sensory modality preferences of grid cells in 2-D spaces (Hafting et al. 2005). Indeed, future studies using 2-D multisensory environments may also be able to determine the properties of the olfactory space map we identified in LEC, where we speculate that grid cells tiling 2D olfactory space may exist.

We found that behavioral demand modulates spatial representations in both LEC and MEC, though to a significantly lesser extent than in CA1 (Figure 5). In the entorhinal cortices, these modulations were largely attributable to non-spatial encoding dimensions, such as speed dependence and reward encoding, which are shaped by the animal’s behavior. Nonetheless, our data suggest that a small fraction of spatial cells in LEC and MEC genuinely encode sensory spaces other than their preferred modalities (see example cells in Figure S1D, S2D, and S3D). Such visual spatial tuning in LEC or olfactory spatial tuning in MEC could arise from feedback projections from the hippocampus. Indeed, the entorhinal–hippocampal network is structured as a recurrent loop, with the hippocampus, via the subiculum, projecting back to both LEC and MEC, modulating their spatial and contextual encoding <u>(Butola et al. 2025; Bonnevie et al. 2013)</u>, and likely contributing to the emergence of some of the cross-modality representations in the entorhinal cortex. For example, LEC receives spatially tuned input from both CA1 <u>(van Strien et al. 2009)</u> and MEC <u>(Nilssen et al. 2019)</u>, and hippocampal feedback activates MEC in response to olfactory stimuli <u>(Biella and de Curtis 2000)</u>. These feedback pathways may underlie the occasional presence of spatial cells encoding non-preferred modalities in LEC and MEC observed here. Alternatively, such cross-modal spatial coding may arise through processing in parahippocampal regions that project to MEC and LEC, including the perirhinal and postrhinal cortices or the claustrum, which have been implicated in multisensory spatial integration <u>(Young et al. 1997; Kitanishi and Matsuo 2017; Connor and Knierim 2017; Nilssen et al. 2019)</u>. For example, LEC receives spatially tuned input from the postrhinal cortex <u>(LaChance et al. 2019)</u>.

Modality-invariant cognition has a neural basis in the hippocampus. For example, single neurons in the human hippocampus and rarely in the entorhinal cortex invariantly represent both the face and written name of an individual <u>(Quiroga et al. 2005)</u>, defining the existence of “concept cells” in the entorhinal-hippocampal network <u>(Quiroga 2012; Quian Quiroga 2023)</u>. Similarly, we found that modality-invariant abstract space mapping appears to be mostly formed in the hippocampus. In the classic model, spatial signals travel through MEC while nonspatial signals pass through LEC, and the two streams converge in the hippocampus to construct a context dependent cognitive map <u>(Eichenbaum et al. 2007; Kerr et al. 2007; Witter et al. 2017; Nilssen et al. 2019)</u>. However, we established that LEC and MEC perform modality-dependent spatial mappings, respectively mapping olfactory and visual spaces and the hippocampus appears to integrate these modality-dependent spatial inputs to build higher level modality-invariant maps for multisensory navigation. These findings provide insight into how abstract representations might be constructed in the hippocampus <u>(Bernardi et al. 2020; Nieh et al. 2021; Samborska et al. 2022)</u>, supporting the idea that the hippocampus is able to select the necessary components to build such spaces from separate, parallel representations in presynaptic regions.

Taken together, our findings support a circuit-level model in which parallel, sensory-modality specific cortical pathways independently construct (relatively task-invariant) spatial maps that are subsequently integrated in the hippocampus into a unified, task-specific multisensory cognitive map. Our results, along with recent studies, support the role of the hippocampus in generating goal-directed spatial representations for action planning <u>(Zutshi et al. 2025)</u>. Indeed, in many cases, hippocampal activity primarily encodes space relative to reward <u>(Sosa et al. 2025)</u>, and its output provides essential information to prefrontal circuits for constructing goal-directed behavioral frameworks <u>(El-Gaby et al. 2024)</u>. Our study suggests that a key to this ability of the hippocampus in generating flexible goal-directed multisensory cognitive maps arises from its inheritance of entorhinal modality-specific spatial maps.

### Limitations of the study

Our study focused specifically on the spatial representations of olfactory and visual modalities, while other sensory modalities were not examined. To minimize external auditory cues, we masked sound using high-volume white noise during experiments. Additionally, mice were not provided with spatially-varying tactile cues for whisker sensing. However, both audition <u>(Rossier et al. 2000; Watanabe and Yoshida 2007; Funamizu et al. 2016; Gao et al. 2020; Nguyen et al. 2024)</u> and touch <u>(Sofroniew et al. 2014; Guyoton et al. 2025)</u> are spatial modalities that rodents use during navigation. Future studies should explore how these senses contribute to spatial representations in diverse multisensory navigation tasks.

We used a linearly increasing and decreasing spatial profile for odorant concentration. However, more ethologically valid odor plume dynamics may influence olfactory spatial representations in the brain <u>(Vickers et al. 2001; Jacobs 2012)</u>. Future studies may further investigate the impact of odor dynamics and spatial profiles on entorhinal-hippocampal maps. Further, during our imaging experiments, we mainly sampled dorsal regions and superficial layers of LEC and MEC. Therefore, our findings about entorhinal function introduced here are not yet fully generalizable to the whole of LEC and MEC regions and their different cortical sublayers.

All imaging experiments in this study were conducted on mice that were well-trained in their corresponding multisensory tasks, and to maximize the number of non-redundant recorded cells, we imaged different planes of a given brain region across days. Given that the LEC is essential for olfactory associative learning, and that suppression of LEC inputs to CA1 impairs olfactory contextual learning <u>(Li et al. 2017)</u>, it would be informative to examine the necessity of LEC in acquiring multisensory navigation tasks—particularly those in which olfaction is a behaviorally relevant and required modality.

It would further be informative to determine whether the same LEC or MEC cells are dedicated to encoding olfactory or visual spaces across tasks. However, in our experimental design, we did not investigate this since we did not train mice to learn to switch between two tasks, e.g., the olfactory-guided task and then the visually-guided task, in one experimental session while imaging the same cells over two tasks. We have previously shown that CA1 spatial cells exhibit global remapping when mice switch between olfactory-guided and visually-guided tasks <u>(Radvansky et al. 2021)</u>, with the sensory spatial mapping identity of cells in the former task not predictive of their identity in the latter task. However, the fraction of CA1 visual and olfactory spatial cells is fully modulated in these two tasks, requiring remapping to occur in the modality-dependent place code of individual cells. In contrast, LEC and MEC preserve their preferred modality-specific mapping across different tasks, thus we predict that the encoding preference of individual cells would be preserved across these tasks. Nevertheless, future work will be needed to further explore this question.

## Supplemental Figures

**Figure S1.**
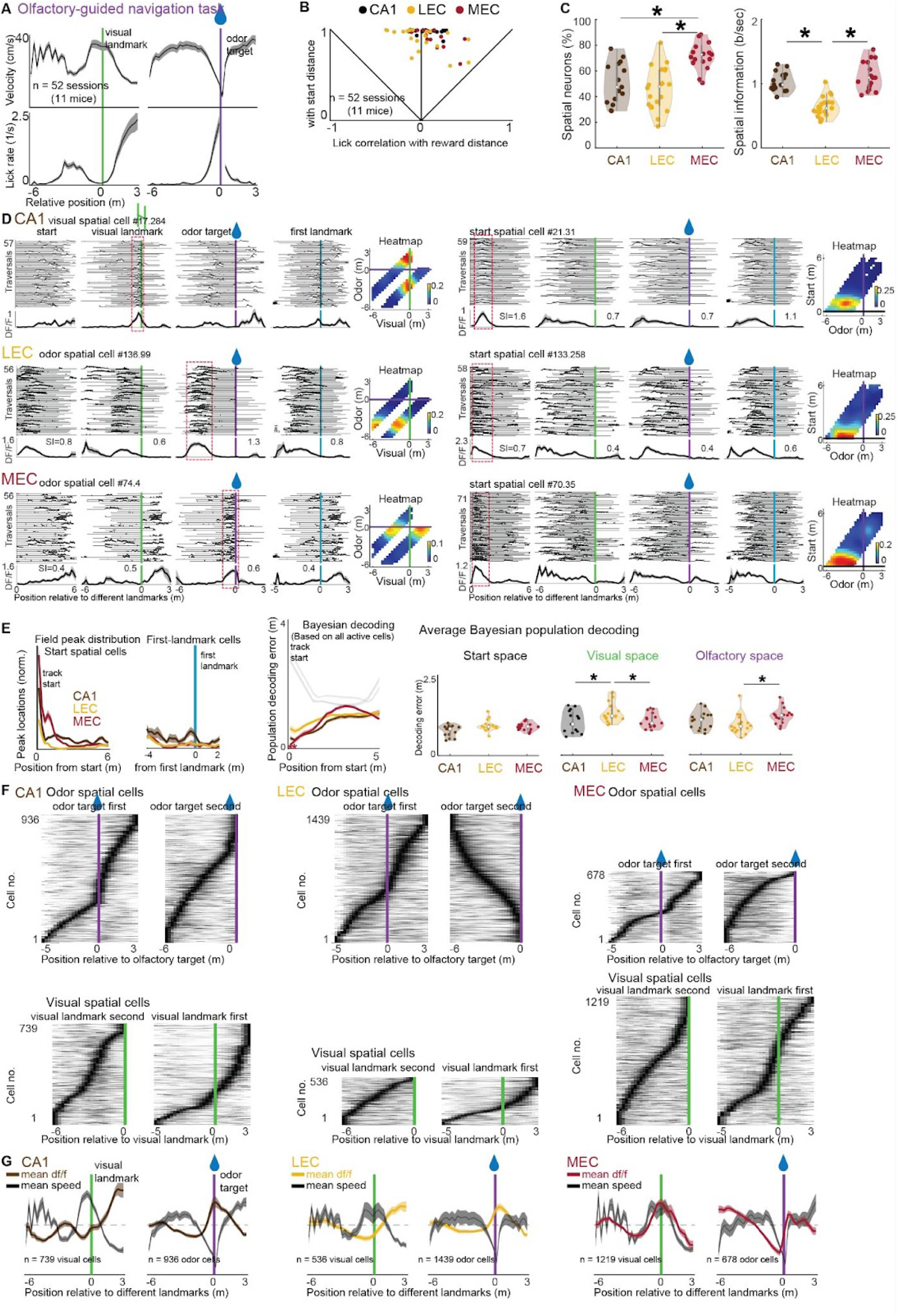
Olfactory-guided navigation task: behavior, spatial cells, and population decoding, related to Figure 2. A) Running and licking behavior of mice in the olfactory-guided navigation task. B) Spearman correlation coefficients between start-to-lick distance and start-to-target distance versus start-to-lick distance and lick-to-target distance plotted for all behavior sessions. Points in the upper quadrant indicate sessions of sensory-guided navigation, in the case of this task, olfactory-guided navigation. Each dot represents a session. C) The proportion of spatial cells among active cells and their spatial information scores in CA1, LEC, and MEC. MEC exhibited the highest fraction of spatial cells relative to CA1 and LEC, with no significant difference observed between CA1 and LEC (two-tailed rank-sum test; CA1 vs LEC, n1 = 15 and n2 = 16 sessions, p = 0.40; CA1 vs MEC, n1 = 15 and n2 = 16 sessions, p = 6.7e-6; LEC vs MEC, n1 = 16 and n2 = 16 sessions, p = 1.1e-5). Additionally, both CA1 and MEC showed significantly higher spatial information scores compared to LEC, with no significant difference between themselves (CA1 vs LEC, n1 = 15 and n2 = 16 sessions, p = 1.3e-5; CA1 vs MEC, n1 = 15 and n2 = 16 sessions, p = 0.54; LEC vs MEC, n1 = 16 and n2 = 16 sessions, p = 6.7e-6). These findings indicate that although LEC has the lowest proportion of spatial cells with the lowest spatial information scores, it nonetheless robustly maps olfactory space, supporting its role in spatial mapping. Each dot represents a session. * p < 0.05. D) Examples of different spatial cell types in CA1, LEC, and MEC. E) Left: Field peak distribution of start spatial cells and Bayesian decoding error of start coordinate based on all active cells. Right: Average Bayesian decoding error of start coordinate, olfactory space, and visual space (two-tailed rank-sum test; Start space: CA1 vs LEC, n1 = 15 and n2 = 17 sessions, p = 0.0541; CA1 vs MEC, n1 = 15 and n2 = 16 sessions, p = 0.0552; LEC vs MEC, n1 = 16 and n2 = 17 sessions, p = 0.90; Visual space: CA1 vs LEC, p = 0.0496; CA1 vs MEC, p = 0.59; LEC vs MEC, p = 0.011; Olfactory space: CA1 vs LEC, p = 0.21; CA1 vs MEC, p = 0.14; LEC vs MEC, p = 0.0065). Each dot represents a session. * p < 0.05. F) Spatial tuning of different spatial cell type populations by separating laps in which a landmark appears first versus second. G) The overlay of mean ΔF/F₀ of different spatial cell types from different brain regions with mice’ average speed.

**Figure S2.**
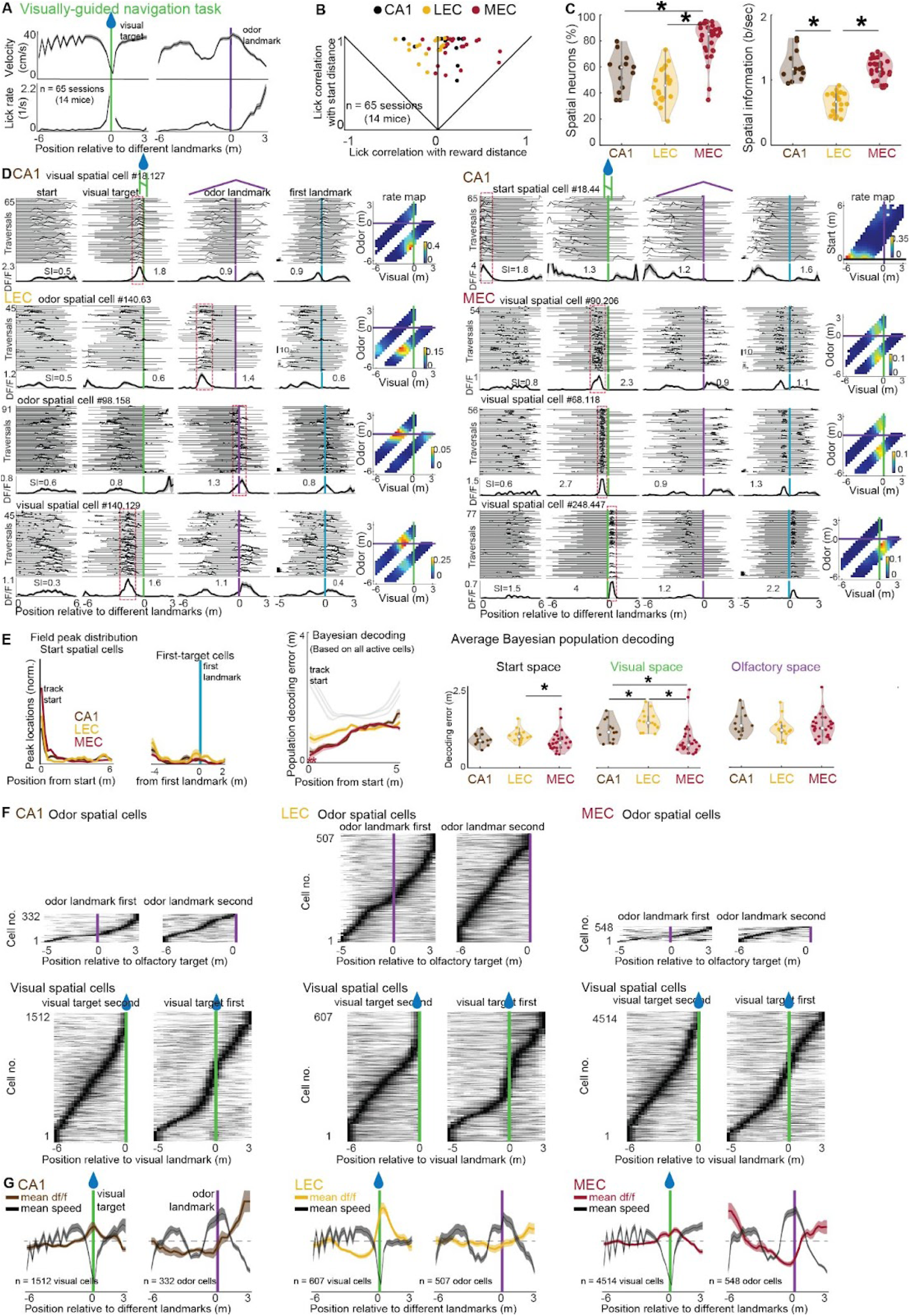
Visually-guided navigation task: behavior, spatial cells, and population decoding, related to Figure 3. A) Running and licking behavior of mice in the visually-guided navigation task. B) Spearman correlation coefficients between start-to-lick distance and start-to-target distance versus start-to-lick distance and lick-to-target distance plotted for all behavior sessions. Points in the upper quadrant indicate sessions of sensory-guided navigation. Each dot represents. C) The proportion of spatial cells among active cells and their spatial information scores in CA1, LEC, and MEC. MEC exhibited the highest fraction of spatial cells relative to CA1 and LEC, with no significant difference observed between CA1 and LEC (two-tailed rank-sum test; CA1 vs LEC, n1 = 13 and n2 = 13 sessions, p = 0.40; CA1 vs MEC, n1 = 13 and n2 = 28 sessions, p = 6.3e-4; LEC vs MEC, n1 = 13 and n2 = 28 sessions, p = 1.1e-5). Additionally, both CA1 and MEC showed significantly higher spatial information scores compared to LEC, with no significant difference between themselves (CA1 vs LEC, n1 = 13 and n2 = 13 sessions, p = 1.7e-5; CA1 vs MEC, n1 = 13 and n2 = 28 sessions, p = 0.77; LEC vs MEC, n1 = 13 and n2 = 28 sessions, p = 6.6e-7). Each dot represents a session. * p < 0.05. D) Examples of different spatial cell types in CA1, LEC, and MEC. E) Left: Field peak distribution of start spatial cells and Bayesian decoding error of start coordinate based on all active cells. Right: Average Bayesian decoding error of start coordinate, olfactory space, and visual space (two-tailed rank-sum test; Start space: CA1 vs LEC, n1 = 13 and n2 = 15 sessions, p = 0.097; CA1 vs MEC, n1 = 13 and n2 = 29 sessions, p = 0.27; LEC vs MEC, n1 = 15 and n2 = 17 sessions, p = 0.0075; Visual space: CA1 vs LEC, p = 0.0304; CA1 vs MEC, p = 0.0023; LEC vs MEC, p = 1.3e-5; Olfactory space: CA1 vs LEC, p = 0.23; CA1 vs MEC, p = 0.43; LEC vs MEC, p = 0.46). Each dot represents a session. * p < 0.05. F) Spatial tuning of different spatial cell type populations by separating laps in which a landmark appears first versus second. G) The overlay of mean ΔF/F₀ of different spatial cell types from different brain regions with mice’ average speed.

**Figure S3.**
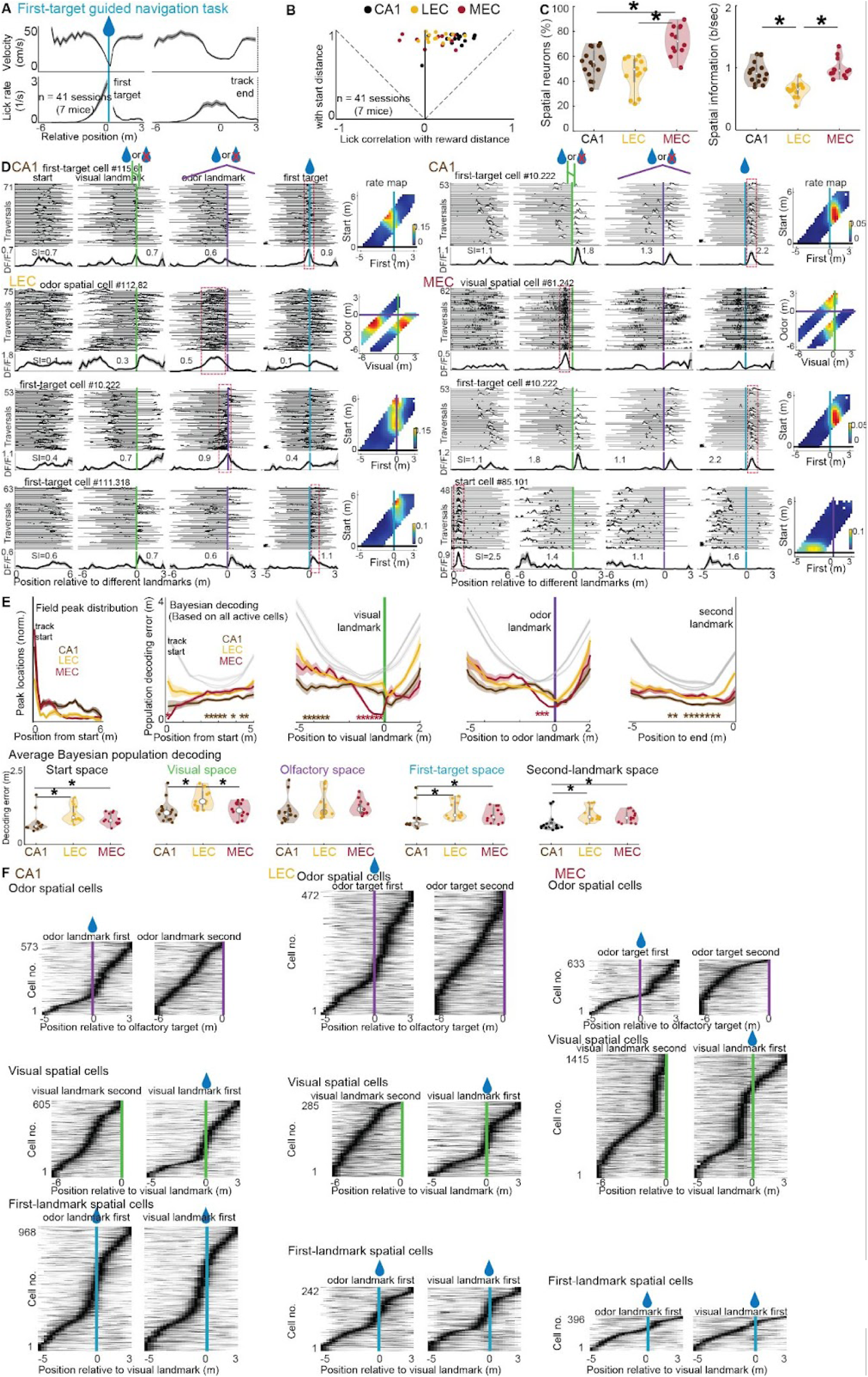
First-target guided navigation task: behavior, spatial cells, and population decoding, related to Figure 4. A) Running and licking behavior of mice in the first-target-guided navigation task. B) Spearman correlation coefficients between start-to-lick distance and start-to-target distance versus start-to-lick distance and lick-to-target distance plotted for all behavior sessions. Points in the upper quadrant indicate sessions of first-target guided navigation. Each dot represents. C) The proportion of spatial cells among active cells and their spatial information scores in CA1, LEC, and MEC. MEC exhibited the highest fraction of spatial cells relative to CA1 and LEC, with no significant difference observed between CA1 and LEC (two-tailed rank-sum test; CA1 vs LEC, n1 = 16 and n2 = 16 sessions, p = 0.44; CA1 vs MEC, n1 = 16 and n2 = 16 sessions, p = 0.0017; LEC vs MEC, n1 = 16 and n2 = 16 sessions, p = 3.2e-4). Additionally, both CA1 and MEC showed significantly higher spatial information scores compared to LEC, with no significant difference between themselves (CA1 vs LEC, n1 = 16 and n2 = 11 sessions, p = 3.8e-5; CA1 vs MEC, n1 = 16 and n2 = 12 sessions, p = 0.32; LEC vs MEC, n1 = 11 and n2 = 12 sessions, p = 5.6e-5). Each dot represents a session. * p < 0.05. D) Examples of different spatial cell types in CA1, LEC, and MEC. E) Left: Field peak distribution of start spatial cells and Bayesian decoding error of start coordinate based on all active cells. Right: Average Bayesian decoding error of start coordinate, olfactory space, and visual space. (two-tailed rank-sum test; Start space: CA1 vs LEC, n1 = 16 and n2 = 13 sessions, p = 0.0006; CA1 vs MEC, n1 = 16 and n2 = 12 sessions, p = 0.0148; LEC vs MEC, n1 = 13 and n2 = 12 sessions, p = 0.101; Visual space: CA1 vs LEC, p = 0.0009; CA1 vs MEC, p = 0.44; LEC vs MEC, p = 0.021; Olfactory space: CA1 vs LEC, p = 0.18; CA1 vs MEC, p = 0.054; LEC vs MEC, p = 0.68; First-target space: CA1 vs LEC, p = 0.0011; CA1 vs MEC, p = 0.0244; LEC vs MEC, p = 0.0971; End coordinate: CA1 vs LEC, p = 0.0008; CA1 vs MEC, p = 0.0043; LEC vs MEC, p = 0.31). Each dot represents a session. * p < 0.05. F) Spatial tuning of different spatial cell type populations by separating laps in which a landmark appears first versus second.

**Figure S4.**
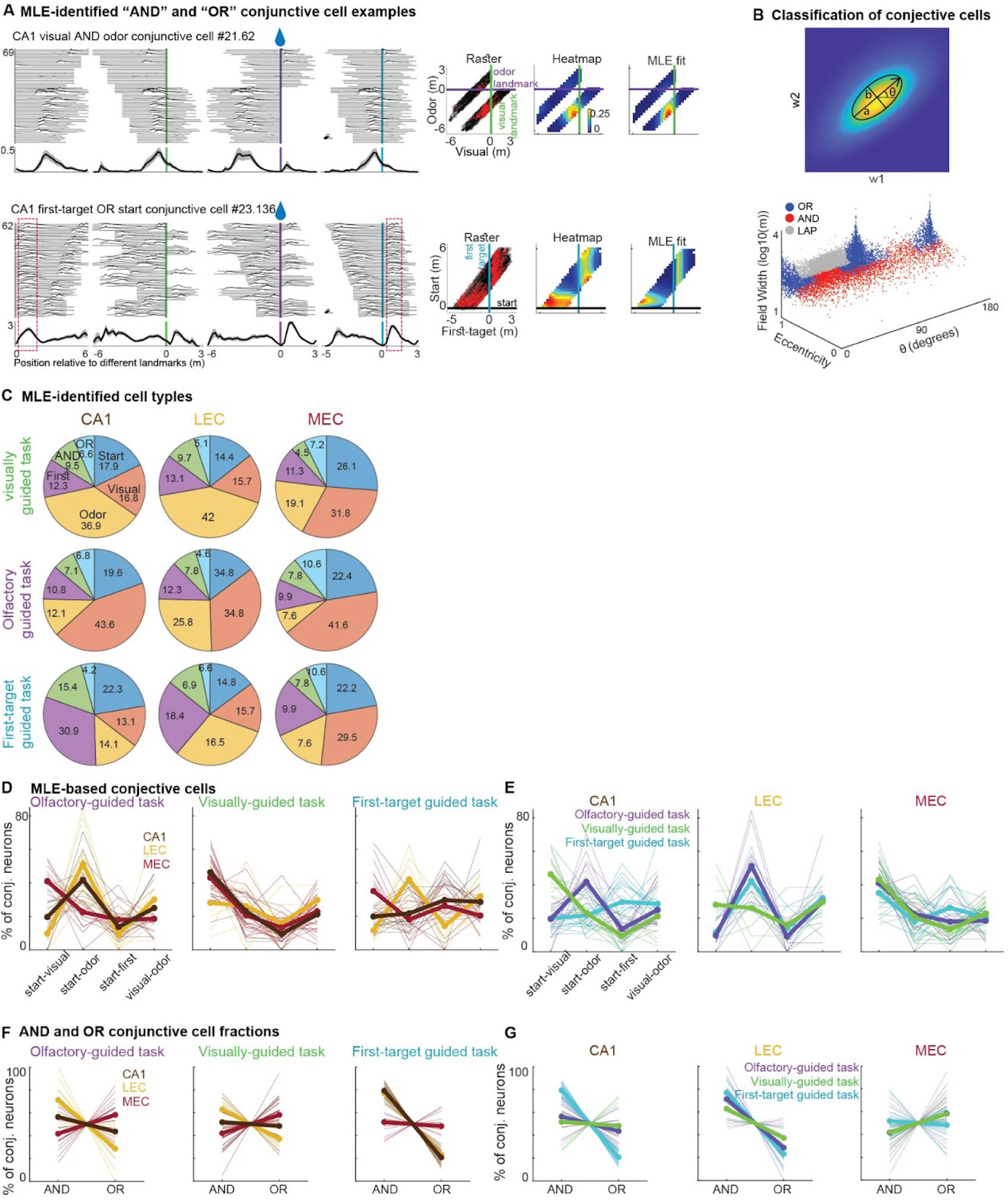
Conjunctive cells: MLE results, related to Figure 5. A) Examples of conjunctive cells in CA1. Top: a visual-odor “AND” conjunctive cell conditionally encode a unique portion of two spaces, i.e., visual and olfactory spaces. Bottom: A start-first target “OR” conjunctive cell independently encodes two spaces, i.e., start and first-target spaces. The two independent start and first-target tuning fields of the OR cell are identified in dashed red rectangles. For each cell, the raster plot of the deconvoluted spikes (red dots) of each example cell is depicted, overlaid on the animal’s position (black dots). The rate map of the raster plot and the fitted MLE model for the example cells are also shown. B) Top: The eccentricity, width, and angle of a hypothetical conjunctive cell is used to classify it as a “AND”, “OR”, or less-selective “LAP” conjunctive cell. Bottom: All MLE-based conjunctive cells are shown and classified based in this three-dimensional classification space. C) The percentage of MLE-based non-conjunctive and conjunctive spatial cells in different multisensory tasks and brain regions. D) The percentage of four MLE-based conjunctive spatial cell types, i.e., start-visual, start-odor, start-first, and visual-odor cell types, in olfactory-guided (left), visually-guided (middle), and first-target guided (right) navigation tasks. Each trace shows a session, with different colors showing different brain regions, and thick colors showing the averages of sessions. Detailed statistics are reported in Table S1. E) The percentage of the same four MLE-based spatial cell types across tasks for each brain region, including CA1 (left), LEC (middle), and MEC (right) recorded during different multisensory tasks. Detailed statistics are reported in Table S1. F) The proportions of AND and OR conjunctive cells in olfactory-guided (left), visually-guided (middle), and first-target guided (right) navigation tasks. Each trace shows a session, with different colors showing different brain regions, and thick colors showing the averages of sessions. Notably, the proportions of OR and AND conjunctive cells differed across hippocampal and entorhinal regions. Detailed statistics are reported in Table S1. G) The proportions of the same AND and OR conjunctive cells across tasks for each brain region, including CA1 (left), LEC (middle), and MEC (right) recorded during different multisensory tasks. While MEC exhibited similar proportions of OR and AND cells, CA1 and LEC contained a significantly higher proportion of AND cells compared to OR cells (detailed statistics in Table S1). These findings suggest that conjunctive cells integrate visual and olfactory spaces through Boolean computations and further highlights the enhanced capacity of CA1, relative to its upstream entorhinal inputs, to perform flexible, modality-invariant computations necessary for multisensory navigation, particularly in the first-target guided task.

**Figure S5.**
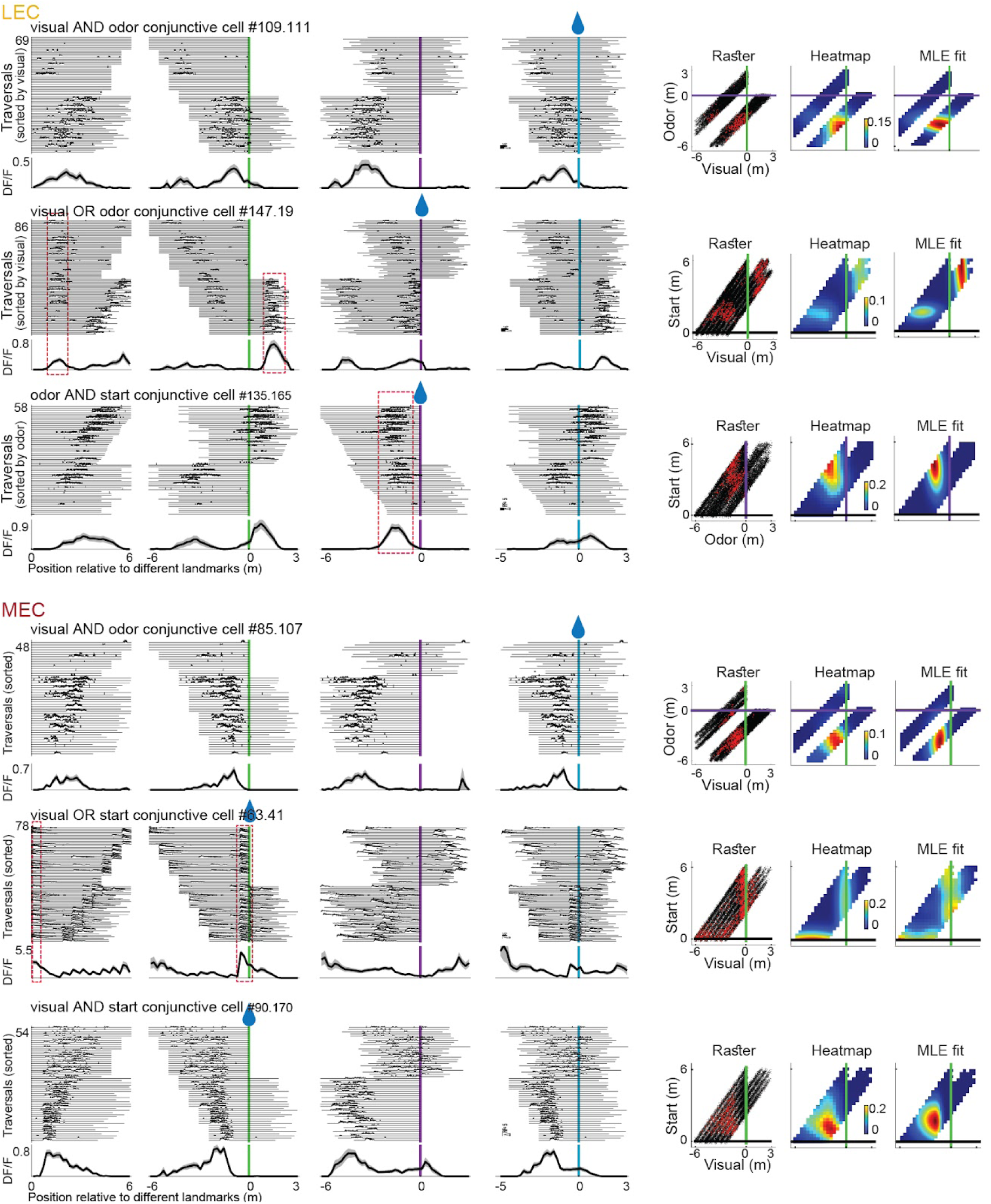
Conjunctive cells: LEC and MEC cell examples, related to Figure 5. Examples of Boolean “AND” and “OR” conjunctive cells in LEC and MEC. “AND” conjunctive cells conditionally encode a unique portion of two spaces, e.g., visual and olfactory spaces. “OR” conjunctive cells independently encode two spaces. The two independent tuning fields of each OR cell are identified in dashed red rectangles. For each cell, the raster plot of the deconvoluted spikes (red dots) of each example cell is depicted, overlaid on the animal’s position (black dots). The rate map of the raster plot and the fitted MLE model for the example cells are also shown.

**Table S1.**
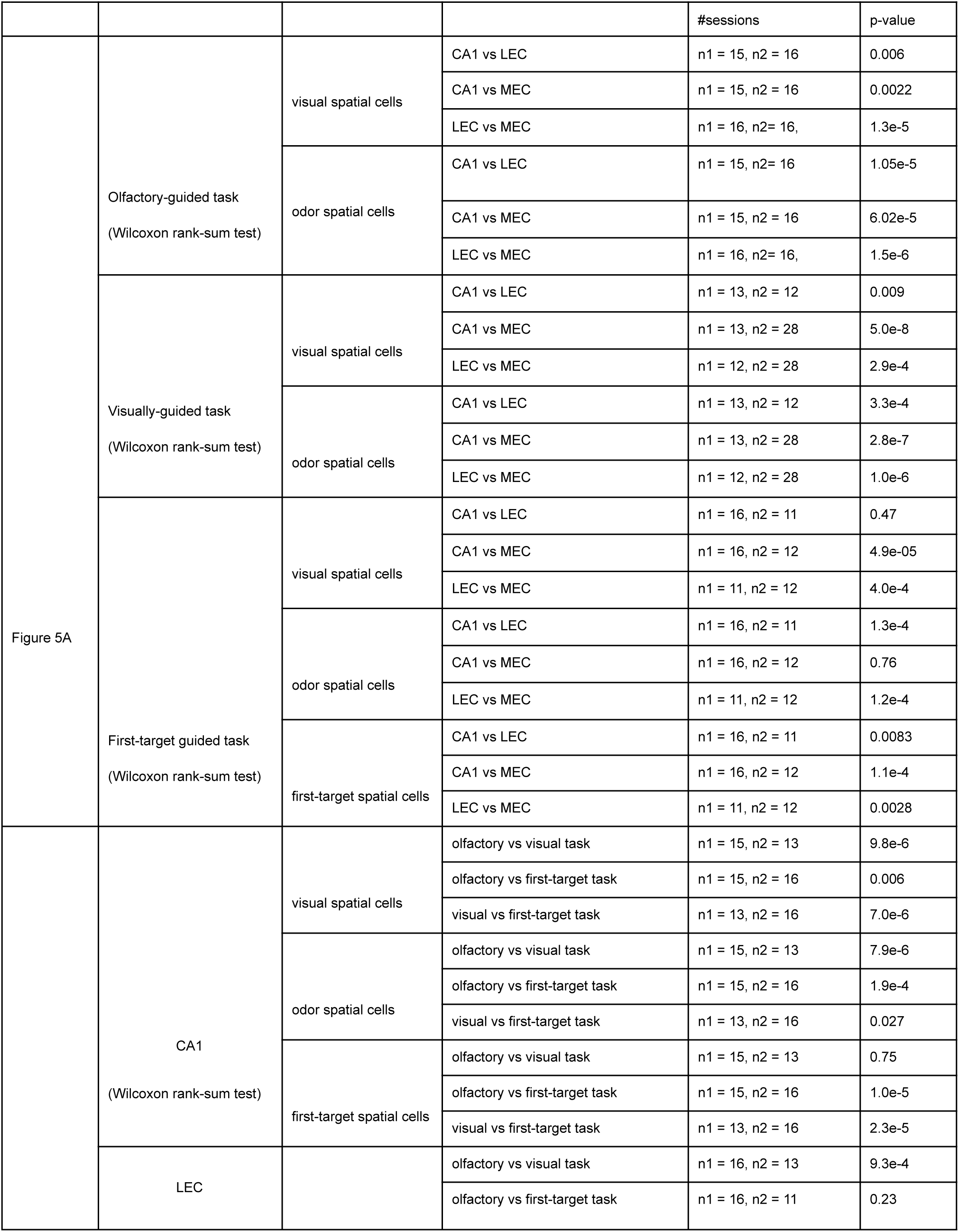
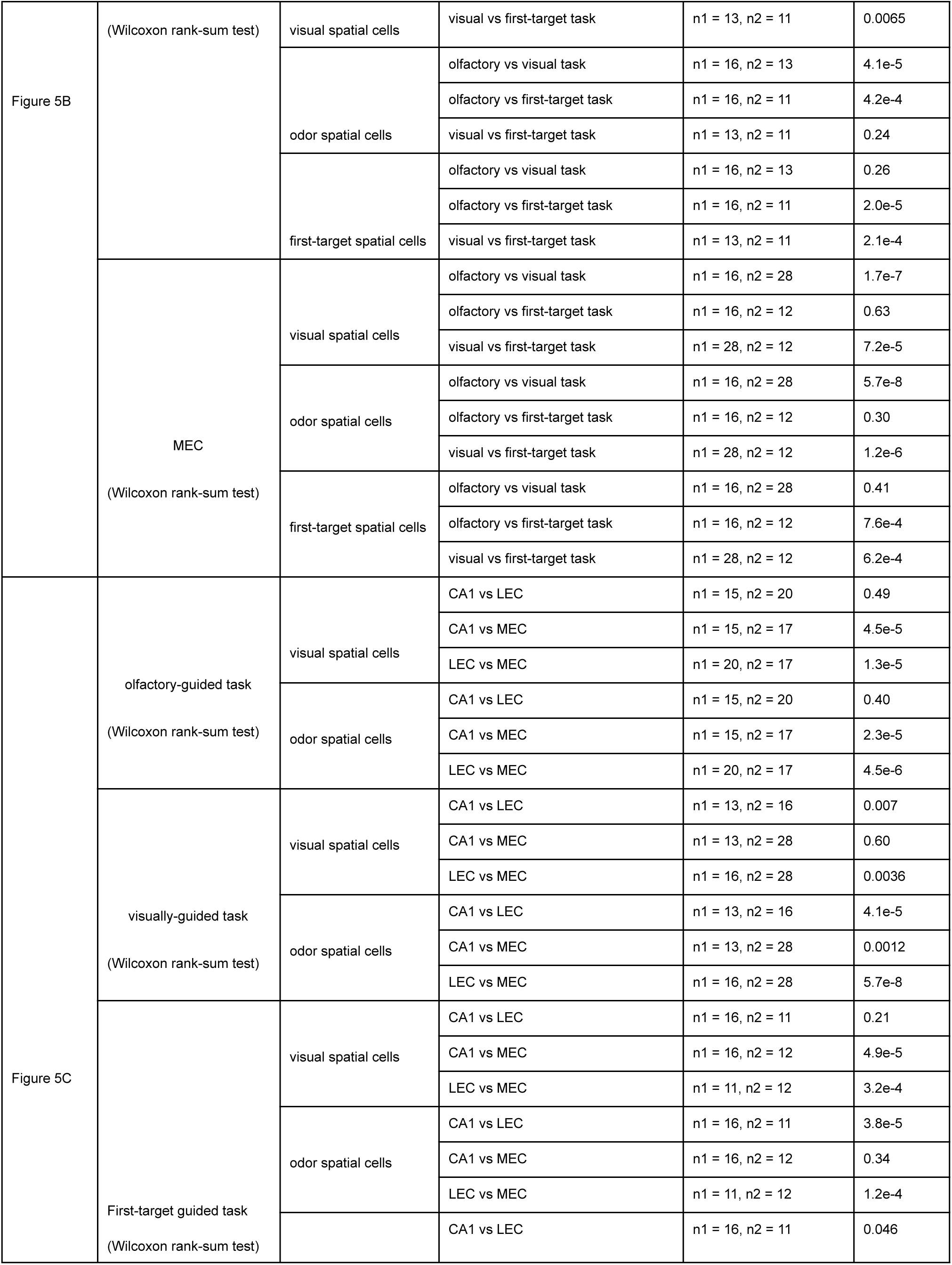
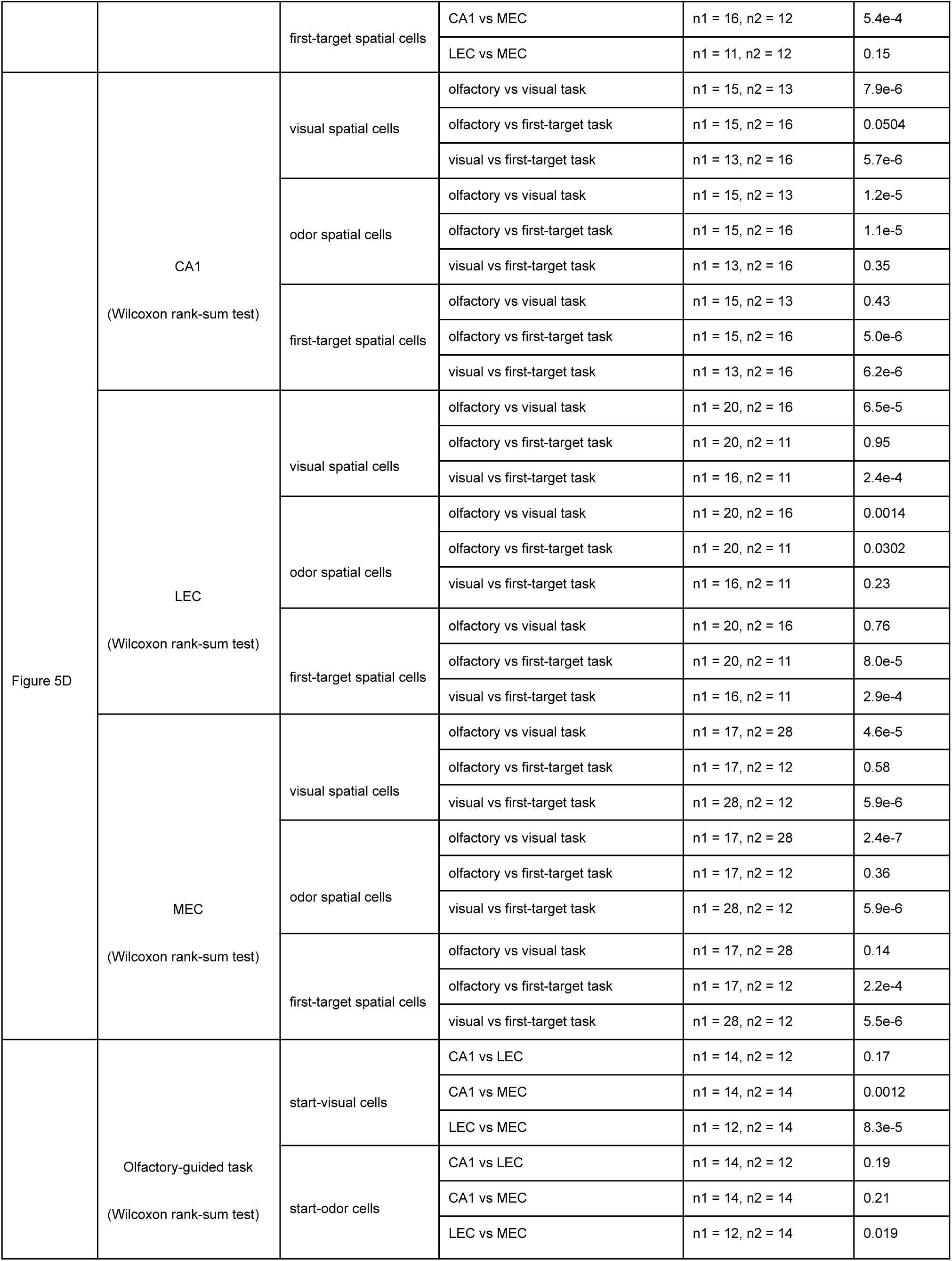
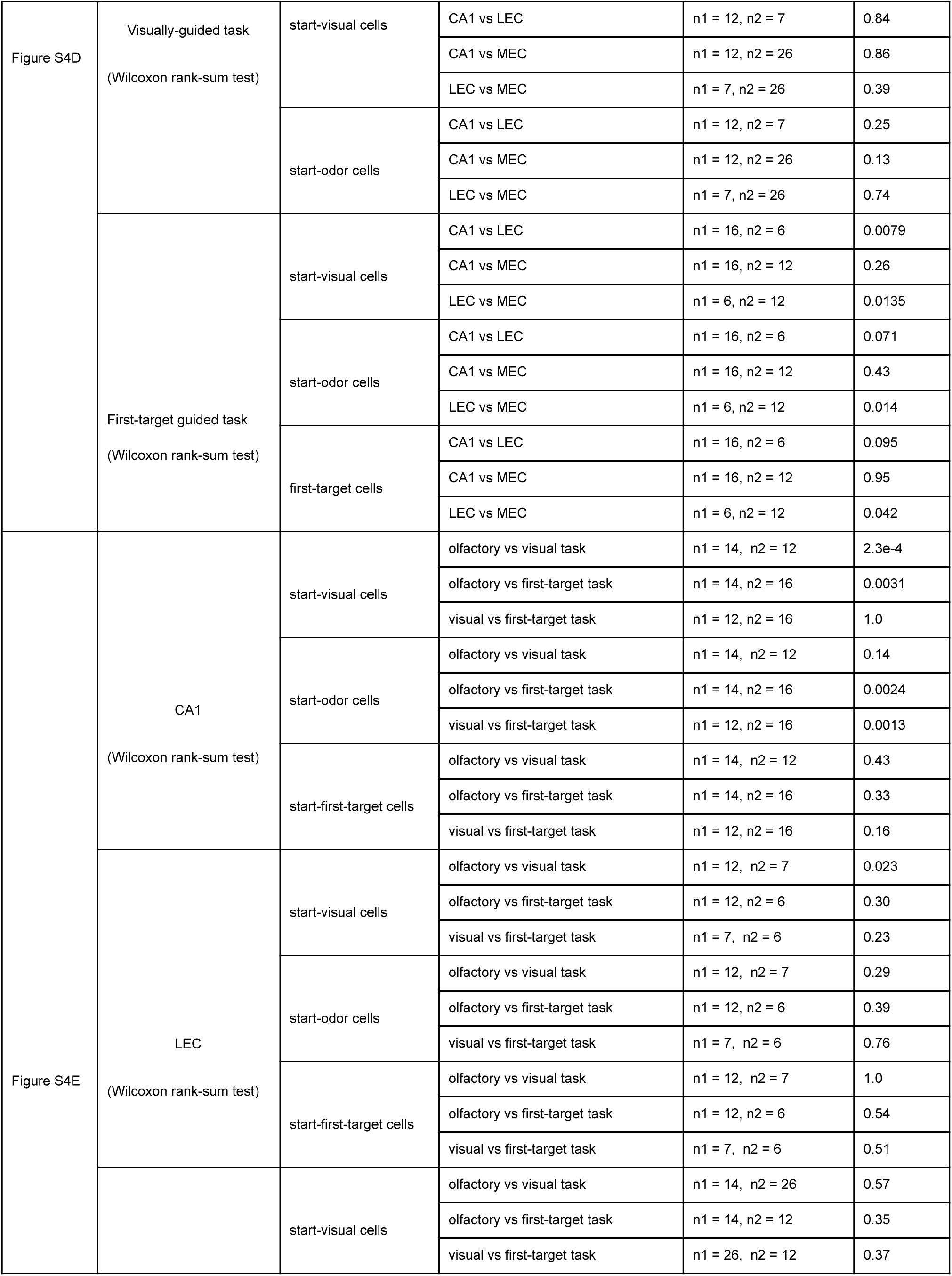
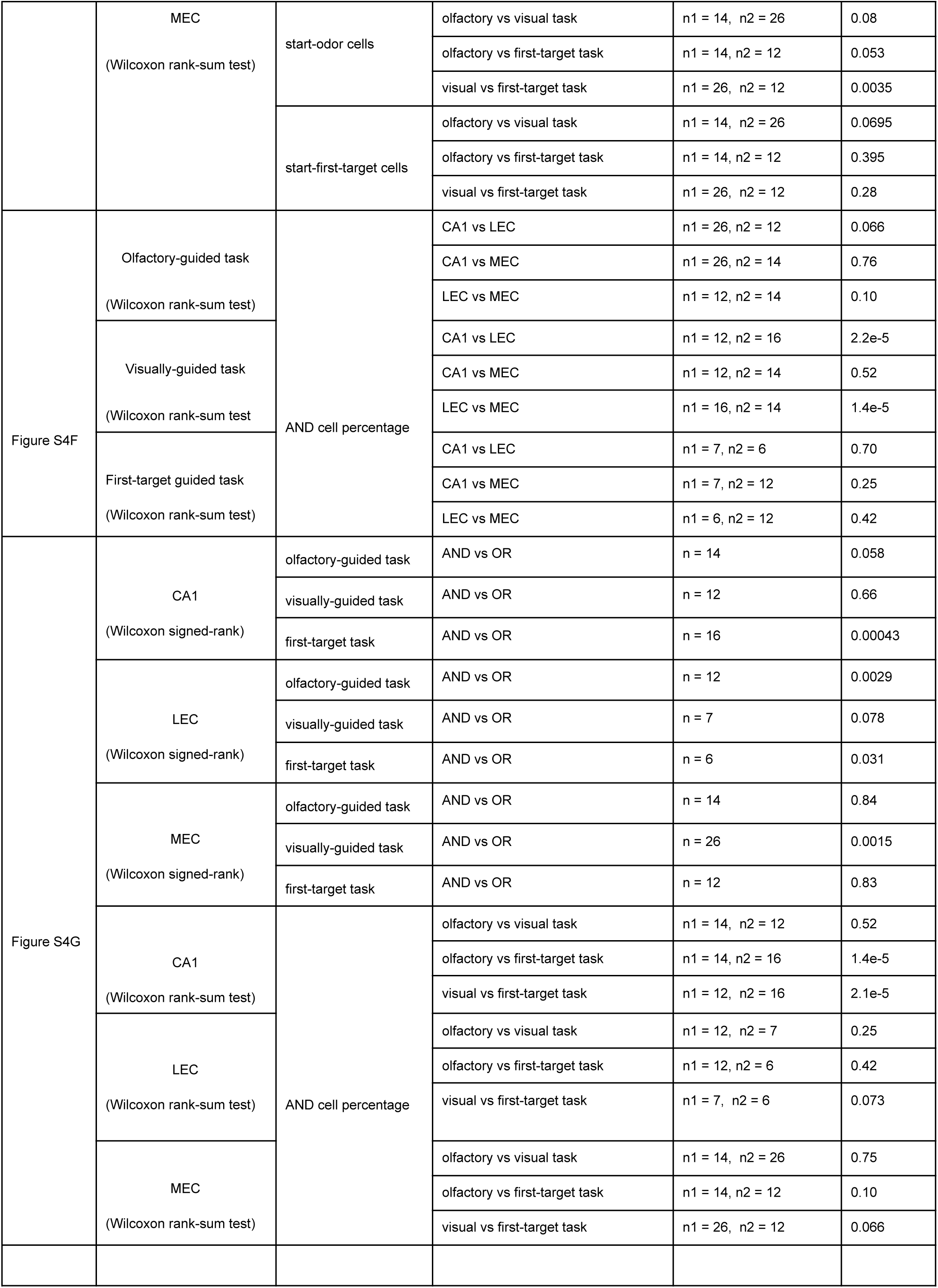
Statistics for task modulation, related to Figure 5 and S4.

## RESOURCE AVAILABILITY

### Lead contact

Daniel Dombeck

### Material, Data and Code availability

All Matlab codes used in this project can be attained upon request. For additional information, or requests pertaining to resources and reagents, please reach out to the lead contact, Daniel Dombeck. All data reported in this paper will be shared by the lead contact upon request. Any additional information required to reanalyze the data reported in this paper is available from the lead contact upon request.

## ACKNOWLEDGEMENTS

This work was supported by NIH grants R01MH101297, T32AG020506, 1F32NS116023 and a NARSAD Young Investigator Grant from the Brain and Behavior Research Foundation (to J.B.I.). The funders had no role in study design, data collection and analysis, decision to publish or preparation of the manuscript.

## STAR METHODS

### KEY RESOURCES TABLE

| REAGENT or RESOURCE | SOURCE | IDENTIFIER |
| --- | --- | --- |
| **Bacterial and virus strains** |  |  |
| pAAV.Syn.GCaMP6f.WPRE.SV40 (AAV1) | Addgene | 100837-AAV1 |
| **Experimental models: Organisms/strains** |  |  |
| tetO-GCaMP6s mice | Jackson Lab. | JAX-024742 |
| Camk2a-tTA mice | Jackson Lab. | JAX-007004 |
| **Software and algorithms** |  |  |
| MATLAB | MathWorks | https://www.mathworks.com |
| ScanImage 5.1 | MBF Bioscience | https://www.mbfbioscience.com/products/scanimage |
| Clampex 10.3 | Molecular Devices | https://www.moleculardevices.com/ |
| Suite2P | Harris Lab | https://github.com/MouseLand/suite2p |
| ViRMEn | Tank Lab | https://github.com/Tank-Lab/ViRMEn |
| FINDBN | Dombeck Lab | https://github.com/DombeckLab/Issa2023 |
| **Deposited data** |  |  |
| Raw and analyzed data, including electrophysiological and behavioral data | This paper | Available upon request |
| Custom code | This paper | Available upon request |

### EXPERIMENTAL MODEL AND STUDY PARTICIPANT DETAILS

All experimental procedures were approved by the Northwestern University Institutional Animal Care and Use Committee. To image neural populations in the LEC and MEC, we bred and used transgenic mice expressing GCaMP6s, generated by crossing tetO-GCaMP6s mice <u>(Wekselblatt et al. 2016)</u> (JAX catalog no. 024742) with Camk2a-tTA mice (JAX catalog no. 007004). For CA1 imaging, these mice received adeno-associated virus (AAV) injections as described below. The number of female/male animals used per region was as follows: CA1 (3M), LEC (2M/4F), and MEC (5M/1F).

### METHOD DETAILS

#### Multisensory Virtual Reality (VR) apparatus

We employed our previously described custom multisensory VR system <u>(Radvansky and Dombeck 2018; Radvansky et al. 2021)</u>, which enables precise and independent manipulation of visual and olfactory cues. Head-fixed mice ran on a cylindrical treadmill positioned in front of a five-panel monitor array. Treadmill velocity was measured using a rotary encoder (US Digital E2-5000) and sampled in MATLAB at a VR engine refresh rate of approximately 5 ms via a National Instruments PCI-6229 data acquisition card. The treadmill rotation signal was scaled to correspond to virtual distance (63.8 cm per revolution). Visual environments were generated in MATLAB using ViRMEn <u>(Aronov and Tank 2014)</u> with cylindrical transformation. The environment consisted of parallel transparent walls featuring white spot patterns to generate optic flow and a single large green arch serving as a spatial visual landmark. The virtual track extended infinitely, with the visual target defined as the crossing of the front plane of the visual landmark.

Olfactory stimuli were delivered using mass flow controllers (MFCs; Alicat MC-100SCCM-D for odorant, flow range 0–0.1 L/min; MC-15SLPM-D for carrier air, flow range 0–1 L/min). The odorant solution (12 mL α-pinene diluted in mineral oil at 1:37.5, Sigma-Aldrich) was housed in a sealed 40 mL vial filled with glass beads and mixed with carrier air in a passive mixing chamber. The resulting odor-air mixture was delivered into a 0.07 cm³ nose chamber positioned around the mouse’s snout. Odor concentration was dynamically modulated (1%–100%) via the odorant MFC, with the carrier stream adjusted in tandem to maintain a constant total flow rate of 1.000 L/min. Along the virtual track, the odor concentration followed a linear spatial gradient with peak centered at the odor landmark (x₀):

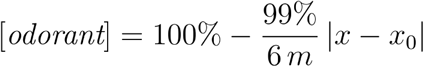

Here, x denotes the mouse’s position, and x₀ marks the location of the odor landmark. Both the gradient slope and the peak concentration were held constant, enabling the mouse to estimate the landmark’s location from any position along the track. To improve spatial accuracy, a predictive algorithm estimated the mouse’s future position 183 ms ahead—compensating for olfactometer delay—and adjusted odor delivery accordingly at each VR iteration.

Licks were detected using a capacitive sensing circuit (SparkFun) connected to a water spout placed within the reach of the mouse’s tongue, and were recorded in MATLAB via a National Instruments PCI-6229 card. All task components—including visual stimuli, MFC control, and reward delivery—were controlled through the ViRMEn engine in Matlab software. A Digidata 1440A system (Molecular Devices; Clampe× 10.3) recorded and synchronized all signals at 1 kHz, including virtual track position, MFC flow rates, licking events, reward delivery, visual and odor landmark encounters, and two-photon imaging frame times.

#### CA1 surgery

Mice were anesthetized with isoflurane (3% for induction, 1–2% for maintenance in 0.5 L/min O₂), with body temperature maintained at 37 °C and ophthalmic ointment applied to prevent corneal drying. To reduce inflammation and manage pain, dexamethasone (5 mg/kg) and buprenorphine-SR-LAB (1 mg/kg, subcutaneous) were administered, along with saline (0.5–1.0 mL) to prevent dehydration. A small craniotomy (∼0.5 mm diameter) was made at coordinates 2.3 mm caudal and 1.8 mm lateral (right hemisphere) relative to Bregma. To express GCaMP6f <u>(T.-W. Chen et al. 2013)</u> in CA1 pyramidal neurons, we injected pAAV.Syn.GCaMP6f.WPRE.SV40 (Addgene #100837-AAV1; diluted ∼10× from a 2 × 10¹³ GC/mL stock in PBS) to a depth of 1.3 mm below the dura using a beveled glass micropipette. Typically, 60 nL of viral solution was injected at the level of the CA1 stratum pyramidale. Two to four days later, a second surgery was performed to implant a stainless-steel cannula with an attached 2.5 mm No. 1 glass coverslip (affixed with NOA81 adhesive, Norland; Potomac Photonics) above the hippocampus <u>(Dombeck et al. 2010)</u>. A titanium headplate was secured to the skull using dental cement (Metabond, Parkell). Mice were monitored postoperatively for 24 h and allowed to recover for 3–5 days before beginning water scheduling (0.9–1.0 mL/day) and behavioral training.

#### LEC surgery

Prism implantation for LEC imaging was performed according to our previously described protocol <u>(Issa et al. 2024)</u>. As the LEC is a lateral region, we used a microprism to rotate the imaging plane by 90°, enabling optical access from above. A 3 mm craniotomy centered 3.5 mm caudal to Bregma was performed over the right lateral skull, positioned as far ventral as possible to cross beyond the rhinal fissure. A 3 mm round No. 0 coverslip (CS-3R-0, Warner Instruments) was temporarily held in place using a micromanipulator-mounted pipette (1.0 mm diameter, Q100-70-7.5, Sutter Instrument) and secured with dental cement (Metabond, Parkell). UV-curable adhesive (NOA81, Norland; UV source: CS20K2, Thorlabs) was then used to affix a 2.0 mm microprism (MPCH-2.0, Tower Optical) to the external surface of the coverslip. Additional dental cement was applied to seal any remaining gaps and to attach a titanium headplate.

#### MEC surgery

Prism implantation for MEC imaging followed our established protocol <u>(Issa et al. 2024; Heys et al. 2014)</u>. A 2–3 mm craniotomy was performed over the right cerebellum. Next, a 2 mm incision was made in the cerebellar dura along the posterior edge of the transverse sinus, and a small amount of cerebellum was aspirated to expose the caudal cortical surface. A 45° microprism (1.5 mm or 2.0 mm; MPCH-1.5 or MPCH-2.0, Tower Optical) mounted onto a custom stainless-steel holder using UV-curable adhesive (NOA81, Norland) was positioned such that its front face contacted the MEC. The prism assembly was then secured with dental cement while applying gentle anterior pressure to enhance stability of the window. A titanium headplate was then attached with dental cement.

#### Behavior

Water-scheduled mice received 0.9–1.0 mL of water per day, with daily monitoring of body weight and overall health. Starting approximately three days after cannula implantation, mice underwent daily training sessions (30–45 minutes) on the multisensory VR system, performing one of three navigation tasks: olfactory-guided, visually-guided, or first-target guided <u>(Radvansky et al. 2021)</u>. In each task, head-fixed mice navigated a one-dimensional multisensory virtual environment by running on a treadmill and received water rewards upon reaching a task-specific target—the odor landmark, the visual landmark, or the first of the two. Mice typically acquired the olfactory-guided and visually-guided tasks within 1–2 weeks, as evidenced by consistent anticipatory licking and slowing behaviors before reaching the target. For the first-target guided task, mice were first trained on the olfactory-guided task for over a week before transitioning to the combined task, reaching criterion performance within an additional 6–9 sessions. Some mice where imaged in multiple tasks.

In each trial, the visual landmark was randomly placed at one of seven possible locations along the track: 2.40, 3.00, 3.60, 4.20, 4.80, 5.40, or 6.00 m from the start. The odor target was positioned randomly at a distance of 1.15–2.99 m either before or after the visual landmark. If the selected position placed the odor target too close to the start (<2 m) or beyond the end of the track (>6 m), it was re-sampled. Total track lengths varied from 3.55 to 6.00 m (mean ± SEM: 5.26 ± 0.64 m), with inter-landmark distances ranging from 1.15 to 2.99 m (2.01 ± 0.48 m). Water rewards (2 µL each) were delivered via a solenoid valve calibrated by its open duration (typically 20 ms). The valve was housed in sound-dampening foam outside the behavioral chamber, and white noise was played during sessions to mask any auditory cues associated with reward delivery. Each trial concluded when the mouse reached the second landmark. At the end of the trial, both visual and olfactory stimuli were frozen for 1 second, followed by a 2-second “washout” period during which the screen turned black and odor concentration was reduced to the minimum (1%). After washout, the mouse was automatically returned to the starting position for the next trial.

#### Two-photon imaging

Imaging sessions began once mice reached stable task performance. Criteria for imaging included: (1) a behavioral rate exceeding two laps per minute on average during 30–50-minute sessions, and (2) consistent anticipatory behaviors across sessions, such as pre-reward licking and slowing. To ensure high-quality behavioral data, imaging in a particular session commenced only after the animal completed several correct trials displaying robust pre-reward licking. All sessions were monitored in real time by the experimenter, and all mice exhibited sufficient anticipatory behavior to be included in further analyses.

Two-photon calcium imaging was conducted using a custom-built, moveable-objective two-photon microscope equipped with a resonant scanner (Sutter Instruments) and a 20X objective (LUCPlanFL N, Olympus). Excitation was provided by a mode-locked Ti:Sapphire laser tuned to 920 nm (Chameleon Ultra II, Coherent), delivering approximately 100-150 mW of average power out of the objective. Emitted fluorescence was separated using a 560 nm longpass dichroic mirror (FF560-Di01, Semrock) into green (FF01-510/84, Semrock) and red (FF01-620/52, Semrock) channels and detected by GaAsP photomultiplier tubes (H10770PA-40, Hamamatsu Photonics). Image acquisition was controlled via ScanImage (Vidrio Technologies). Time-series recordings typically consisted of 16,000 frames at 512 × 256 pixels (0.0675 ms/line), acquired at 58.4 Hz. Most sessions ranged from 64,000 to 96,000 total frames. For LEC and MEC prism imaging, the field of view ranged from 300 × 300 µm to 600 × 600 µm, with imaging typically performed at 1.1X to 1.9X magnification.

To minimize contamination from the VR monitor’s light, a custom light-shielding cylinder made from opaque electrical tape was positioned between the headbar and the microscope objective.

### QUANTIFICATION AND STATISTICAL ANALYSIS

#### Statistics and reproducibility

All data analyses were performed using custom-written software in MATLAB (MathWorks). Although statistical methods were not used to predetermine sample sizes, the number of subjects was consistent with prior publications <u>(Kaufman et al. 2020; Radvansky et al. 2021; Issa et al. 2024)</u>. The experimenter was not blinded to experimental conditions during data collection or analysis due to the necessity of anatomical targeting for imaging, which made blinding impractical. Inclusion criteria were based on predefined behavioral performance metrics, and all mice with high-quality imaging windows and satisfactory task performance were included in the analysis.

Only non-parametric statistical tests were used, with no assumptions about underlying data distributions. These included bootstrapping, Wilcoxon signed-rank tests, Wilcoxon rank-sum tests, and chi-square goodness-of-fit tests, applied where appropriate. For multiple comparisons using the Wilcoxon rank-sum test, uncorrected p-values are reported, with Bonferroni-corrected thresholds for statistical significance indicated in figure legends. Statistical significance was defined as p < 0.05, except for neuron-type classification analyses, where a stricter criterion of p < 0.01 was applied. While data distributions were assumed to be approximately normal, no formal tests for normality were conducted. All quantitative results are reported in the main text and figure legends, including definitions of sample size (n), measures of central tendency, and statistical significance thresholds. Unless otherwise specified, values are reported as mean ± SEM. Cross-validation was performed by comparing results from odd and even trials, and analyses were conducted both at the session level and across animals to assess generalizability. Imaging experiments across brain regions were conducted using identical experimental protocols and analysis pipelines. Regional comparisons were not randomized, as recordings for each area were typically carried out in dedicated experimental batches.

#### Behavioral analysis

At each time point, the mouse occupied a single position, defined as its distance from the start of the virtual track. To align behavioral and neural data with task-relevant landmarks, we calculated relative position vectors by subtracting the location of each landmark—start, visual landmark, odor landmark, first landmark, or second landmark—from the current position. Our primary behavioral metric was pre-target licking, used to quantify anticipatory behavior. To isolate anticipatory licks from consummatory licks, we excluded all licks that occurred after reward delivery, within 0.25 m beyond the reward site, or within 0.25 m of the start position. Behavioral sessions were included only if they met two criteria:

1. Target-selective licking—increased lick rates prior to the rewarded target compared to the non-rewarded landmark. For each session, we divided the track into 25-cm bins centered on each landmark and computed lick rates as the number of licks divided by the time spent in each bin. Pre-target licking was defined as the lick rate in the 25-cm bin immediately before the rewarded landmark. Sessions were included if mice exhibited significantly greater pre-target licking for the rewarded versus non-rewarded target across trials, assessed using Wilcoxon signed-rank tests.
2. Navigation strategy—to distinguish sensory-guided navigation from path integration, we assessed whether mice adjusted the location of their licking based on the landmark position. For each trial, we computed the median lick position occurring between the start and the rewarded target. We then plotted this median position in both start-aligned and target-aligned coordinates against the distance between start and target. Spearman correlation coefficients were calculated: R-start (correlation in start-aligned coordinates) and R-target (correlation in target-aligned coordinates). Sessions were classified as sensory-guided if R-start > R-target, indicating lick position was dependent on the location of the target rather than a fixed distance from the start. For the first-target guided task, this analysis was performed separately for visual-first and odor-first trials to confirm strategy consistency across modalities.

Only sessions that met both criteria—pre-target licking and non-path-integrative navigation—were included in the final dataset.

#### Calcium Imaging analysis

We applied the following analysis pipeline using custom MATLAB code. First, motion correction, ROI segmentation, and raw fluorescence extraction were performed using a Python implementation of Suite2p <u>(Pachitariu et al. 2017)</u>. For each ROI, we extracted the raw fluorescence vs time trace along with a corresponding neuropil signal. A small fraction of traces were removed due to out of range min, max, and mean values. We then applied an integrated, iterative algorithm <u>(Issa et al. 2024)</u> to calculate ΔF/F₀ by estimating the neuropil contamination ratio and baseline fluorescence (F₀). Next, significant calcium transients were calculated from ΔF/F₀ <u>(Dombeck et al. 2007)</u> and this signal was used for the subsequent calculations throughout the papers.

Behavioral data were downsampled to match the imaging frame rate (58.4 Hz) and aligned with the timing of significant calcium transients. To isolate activity related to active navigation, we restricted the analysis to time points when the animal was making forward progress along the track (> 10 cm/s). Only movement-associated calcium events were included in subsequent neural analyses, excluding periods of rest, reward consumption, or inter-trial intervals.

#### Spatial information calculation and spatial cell type classification

To identify active neurons, we first computed the average ΔF/F₀ signal for each cell across all laps and time points within a session. Neurons were classified as “active” if they contained significant calcium transients on at least 1/3 of laps. Imaging sessions that contained more than 50 active cells were considered for further analysis. Spatial information of each individual cell was calculated using our previously established method <u>(Climer and Dombeck 2021; Radvansky et al. 2021)</u>. Only imaging frames corresponding to running periods (> 10 cm/s) were included in the analysis. Spatial information content was computed for each neuron using the following formula:

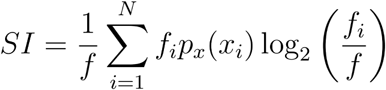

Where *SI* is the spatial information in bits per event, *N* is the total number of spatial bins (60 bins, bin width = 25 cm), *f_i_*is the mean fluorescence (inferred firing) in bin *i*, *f* is the overall mean fluorescence across all bins, and *p_x_(x_i_)* is the occupancy probability of bin *i*. This calculation was performed independently for each spatial reference frame: position relative to the start, the visual landmark, the odor landmark, and the first landmark. As a control, spatial information was also calculated relative to the second landmark in first-target guided navigation sessions.

To classify neurons as start-, odor-, visual-, or first-target–spatial, we adapted a method based on previous approaches <u>(Gothard et al. 1996; Radvansky et al. 2021)</u>. For each neuron, we identified the spatial reference frame (e.g., start, visual, olfactory, or first target) in which the neuron exhibited the highest spatial information score. To determine whether this tuning exceeded chance, we generated a null distribution by shuffling the neuron’s significant transient trace. The trace was divided into ≥ 100 segments (preserving the shape and timing of individual transients), randomly reordered, and then recombined. Spatial information was then recalculated using the original position vector. This process was repeated 1,000 times. The p-value was defined as the proportion of shuffled information values that exceeded the actual observed score. Neurons with p < 0.01 in their highest-scoring spatial frame were classified as spatially tuned to that coordinate. Neurons that met this criterion were classified as “spatial cells”, with no minimum spatial information threshold used. In contrast, neurons that did not meet this threshold were labeled “none.” Although some neurons exhibited significant spatial information in multiple frames, we did not classify them as conjunctive (e.g., olfactory-visual) because the spatial reference frames were not fully independent. For example, strong tuning to the visual landmark could artificially elevate spatial information in the odor coordinate due to the fixed inter-landmark distance (1.15–2.99 m), rather than reflecting true multimodal selectivity. For a better visualization of population heatmaps, the spatial tuning of individual spatial cells were smoothed using a sliding window of 3 bins (each bin width = 25 cm).

#### Bayesian reconstruction

To assess the accuracy of spatial representations across CA1, LEC, and MEC, we used Bayesian decoding methods <u>(K. Zhang et al. 1998; Etter et al. 2020; Radvansky et al. 2021)</u>. The virtual track was divided into 10 cm bins, and data were split into training (odd-numbered trials) and testing (even-numbered trials) sets. For single-trial decoding, we applied leave-one-out cross-validation, training the decoder on all but one trial and evaluating performance on the held-out trial. Each session was divided into time bins of Δt = 0.1 s.For each time bin n, we estimated the conditional probability that the animal was at position bin *x_i_* (width = 25 cm) given the observed significant transients, using the following formulation:

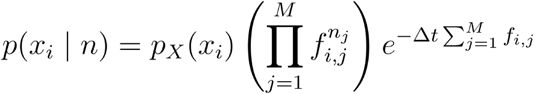

Where *p*(*x_i_* | *n*) is the occupancy probability of bin *x_i_, f_i,j_* is the average transient rate of neuron j in bin *x_i_, n_j_* is the number of transients from neuron j in time bin n, M is the total number of neurons. The decoded position was taken as the bin with the highest posterior probability. Decoding was performed separately using four coordinate systems: distance from the start, from the visual landmark, from the odor landmark, and from the first landmark. As a control, decoding was also performed using distance from the second landmark in the first-target guided navigation task. To establish a chance-level baseline, decoded positions were randomly shuffled across time bins within each trial. Reconstruction error was computed as the Euclidean distance between the real (non-binned) and decoded positions at each time point. The median decoding error within each spatial bin along the track was used due to its robustness against outliers, particularly at the track edges where decoding variability tends to increase. This procedure was repeated for both the original and shuffled data. To compare decoding accuracy across navigation conditions, we used Wilcoxon rank-sum tests applied separately within each spatial bin. A decoding condition was considered significantly more accurate in a given bin if its median error was lower than that of all other conditions (p < 0.05).

#### Maximum likelihood estimation (MLE) for conjunctivity analysis

To identify conjunctive spatial tuning beyond what could arise from overlapping coordinate systems (e.g., start, visual, odor, and first-target coordinates), we employed an MLE framework <u>(Brown et al. 2003; Radvansky et al. 2021)</u> , modeling each neuron’s spiking activity as an inhomogeneous Poisson process. First, calcium traces (ΔF/F₀) from each neuron—including both significant and non-significant transients—were deconvolved using a first-order autoregressive model (AR(1)) with a hard threshold (ℓ₀ penalty) to estimate continuous activity <u>(Friedrich et al. 2017)</u>. Noise levels and the decay constant were initialized manually (τ = 0.58 s). These deconvolved traces were converted into pseudo-spike counts by identifying events that coincided with both a significant transient and a deconvolution peak. A unit pseudo-spike was defined as the median amplitude of the lowest 5% of such events; if insufficient events were present, the smallest was used. All deconvolved values were linearly scaled to this unit and rounded to the nearest integer to yield frame-wise pseudo-spike counts. While this transformation does not claim to capture true spike rates precisely, it provides a reliable and unbiased estimate of relative spiking, allowing consistent comparisons across spatial reference frames within each neuron.

Each neuron’s pseudo-spike rate was modeled as the sum of a skewed multivariate Gaussian tuning function for each field (Azzalini 2005):

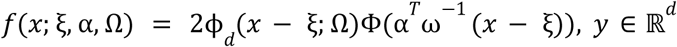

Where ξ is a *d* dimensional location parameter, α is a *d* dimensional shape parameter, Ω is a positive definite matrix, ϕ is the density function of a *N_d_* (0, Ω) variable, and Φ is the cumulative distribution function of a *N*(0, 1) variable.

The covariance matrix of the distribution is 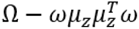 and the skewness is 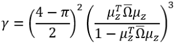 , Where 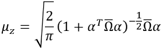 and 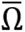 is the correlation matrix associated with Ω. Twice the square root of the eigenvalues of the covariance matrix were used as the widths of the field along its major and minor axes and the eigenvectors were used to determine the field orientation.

Each neuron’s pseudo-spike rate was modeled with the conditional intensity function:

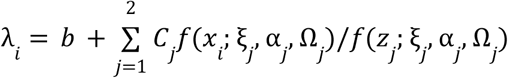

Where *i* is the time bin, *b* is the baseline rate, *j* is the field number, *C* is the peak rate above baseline for the field, *x* is the position of the animal, and *z_j_* is the mode of the *j*th field (determined numerically).

The log-likelihood was determined using a Poisson model:

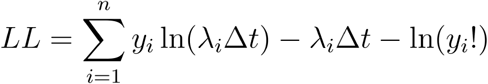

where n is the number of imaging frames, *y_i_* is the observed pseudo-spike count at frame i, and Δt is the frame duration (0.01728 s).

The null model was “flat” (*C*_1_ = *C*_2_ = 0). We fit up to two-dimensional conjunctions (start-visual, start-odor, start-first, and odor-visual) with the exception of the visual-first and odor-first conjunctions due to redundancy in the definition of these coordinates. For “pure” models (start, visual, odor and first), *d*=1, and for “conjunctive” models *d*=2. For each pure and conjunctive dataset, a single field model (*C*_1_ > 0, *C*_2_ = 0) and a two-field model (*C*_1_ > 0, *C*_2_ > 0) was fit.

Altogether, 17 models were fit for each neuron. All parameters were optimized using MATLAB’s particleswarm algorithm. The solutions were constrained so that the minor width of the field was between 20 and 200 cm, the peak of the distribution was at most one standard deviations away from occupied locations in the *d* dimensional conjunctive space, at least ⅓ of the spikes were predicted to fall within each field, and the skewness was at most 0.8, the rate on the line between the field centers fell to at least 10% of the peak.

To select the best-fitting model, each model was fit using five-fold cross validation across trials stratified across which landmark occurred first. The best model was selected using the modified one-standard error rule on the resulting loglikelihood <u>(Yates et al. 2021)</u> and was used to classify the neurons. The MLE approach resulted in eight distinct cell types (i.e., start, visual, odor, first-target, start-visual, start-odor, start-first, and visual-odor spatial cells). The full (non-cross validated) dataset was then fit with the best-fitting model for further analysis. Sessions that contained more than thirty MLE-identified spatial cells were considered for prevalence analysis across tasks and brain regions.

“AND” and “OR” conjunctive cell identification

To further characterize conjunctive neurons, we analyzed their tuning geometry across regions and task conditions, revealing distinct classes of Boolean-like spatial coding. Based on MLE fits and deconvolved spiking heatmaps, we categorized fields from conjunctive neurons into three types using the field widths and orientations.

1. “AND” Fields: These fields encoded combinations of spatial coordinates in an interdependent manner, responding only when both spatial dimensions (e.g., visual *and* odor) were simultaneously satisfied. These were all fields that did not qualify as “LAP” or “OR” fields.
2. “OR” Fields: These fields strongly encoded a single coordinate, as indicated by appearing as a stripe across the headmap.The major axis was within 30 degrees of the cardinal axis, the eccentricity 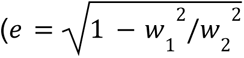, where *w*_1_ is the length of the minor axis and *w*_2_ is the length of the major axis) was at least 0.8, and the major field axis was at least 100 cm.
3. “LAP” Fields: These less-selective fields aligned to the animals motion through the conjunctive space and reflected activity across full laps. They had an eccentricity of at least 0.8, a major field width of at least 300 cm, and were oriented within 30 degrees of the 1:1 line.

A cell was determined to be an AND cell if all its fields were AND type and an OR cell if at least one of its fields were an OR type. LAP cells were broadly tuned and thus excluded from further conjunctive analysis. Sessions that contained more than a total of ten AND and OR cells were considered for prevalence analysis across tasks and brain regions.

